# Parameter-Efficient Fine-Tuning Enhances Adaptation of Single Cell Large Language Model for Cell Type Identification

**DOI:** 10.1101/2024.01.27.577455

**Authors:** Fei He, Ruixin Fei, Mingyue Gao, Li Su, Xinyu Zhang, Dong Xu

**Affiliations:** Department of Electrical Engineering and Computer Science, Bond Life Sciences Center, University of Missouri, Columbia, MO, 65211, USA; School of Information Science and Technology, Northeast Normal University, Changchun Jilin 130017, China

**Keywords:** Single cell, Foundation model, PEFT, Cell type identification

## Abstract

Single-cell sequencing transformed biology and medicine, providing an unprecedented high-resolution view at the cellular level. However, the vast variability inherent in single-cell sequencing data impedes its utility for in-depth downstream analysis. Inspired by the foundation models in natural language processing, recent advancements have led to the development of single-cell Large Language Models (scLLMs). These models are designed to discern universal patterns across diverse single-cell datasets, thereby enhancing the signal-to-noise ratio. Despite their potential, multiple studies indicate existing scLLMs do not perform well in zero-short settings, highlighting a pressing need for more effective adaptation techniques. This research proposes several adaptation techniques for scLLMs by preserving the original model parameters while selectively updating newly introduced tensors. This approach aims to overcome the limitations associated with traditional fine-tuning practices, such as catastrophic forgetting and computational inefficiencies. We introduce two Parameter-Efficient Fine-Tuning (PEFT) strategies specifically tailored to refine scLLMs for cell type identification. Our investigations utilizing scGPT demonstrate that PEFT can enhance performance, with the added benefit of up to a 90% reduction in parameter training compared to conventional fine-tuning methodologies. This work paves the way for a new direction in leveraging single-cell models with greater efficiency and efficacy in single-cell biology.

## Introduction

Single-cell sequencing has significantly advanced the fields of biology and medicine by providing high-resolution insights at the cellular level. This technology offers valuable understanding of the roles and relationships of different cell types within their native environments, shedding light on complex tissues and biological systems where cell-to-cell variation plays a critical role[1]. Diseases such as cancer involve subsets of cells that diverge genetically and behaviorally from normal cells. Single-cell sequencing has the capacity to reveal these subtle yet crucial differences, offering a detailed view of the cellular composition of tumors or the diversity of immune cells in response to infection or treatment, thereby paving the way for personalized medicine[2]. However, single-cell sequencing is accompanied by several technical challenges and limitations, including batch effects[3], uneven coverage[4], dropout[5], potential cellular damage[6], and the introduction of bias and artifacts in the data. These factors complicate downstream analyses and interpretation.

The success of foundation models in natural language processing (NLP) and computer vision (CV)[7][8] provides strong evidence that foundation models can capture universal patterns in data for various downstream analyses. These models boast a massive number of parameters and are pretrained on large datasets, allowing them to focus on understanding broad regularities in the data rather than any specific end-task. The patterns they capture often represent high-level features common across different data types, enabling easy transfer to specific domains and resulting in improved performance on multiple tasks compared to task-specific models trained from scratch[9]. Motivated by these benefits, emerging research has begun to explore the potential of foundation models in single-cell biology, particularly in single-cell transcriptomics. This includes models such as scBERT[10], Genefomer[11], scGPT[12], scFoundation[13], SCimilarity[14], GeneCompass[15], and scTab[16], which aim to pretrain foundation models on large-scale single-cell atlases to yield universal patterns that embed the biological essence of single-cell data and overcome technical issues.

These models, collectively referred to as single-cell large language models (scLLMs), have attracted significant attention and subsequent research[17-24], which has investigated their reusability, extendibility, and applicability. For example, Kasia Z. Kedzierska et al.[17] benchmarked scGPT and Geneformer in zero-shot settings and found that these models did not perform well in such scenarios. Similarly, Boiarsky et al. observed similar results when benchmarking scGPT and scBERT[18]. These findings indicate that the current scLLMs have not yet demonstrated emerging intelligence. Therefore, adapting current scLLMs is crucial for complex tasks, such as cell type identification and gene expression prediction. When Alsabbagh et al.[19] and Kedzierska et al. finetuned scGPT and Geneformer on a small amount of additional training data for benchmarking, they both observed that scGPT outperformed Geneformer in cell type annotation. However, Liu et al. reached the opposite conclusion[20] when finetuning scGPT and Geneformer. These inconsistent results from the same scLLMs highlight the importance of a proper adaptation approach for maximizing the benefits of scLLMs.

The adaptation approaches for scLLMs have been relatively underexplored, with existing works predominantly relying on the traditional fine-tuning approach. However, traditional fine-tuning of large models can lead to the overwriting of original model parameters on narrow, task-specific datasets, potentially resulting in the loss of broader pre-learned knowledge and the phenomenon known as catastrophic forgetting. This, in turn, can lead to a reduction in adaptability and an increased risk of overfitting to the limited training data. Additionally, the resource-intensive nature of fine-tuning large models further exacerbates these challenges. In this study, our primary hypothesis is that a more effective adaptation approach for scLLMs involves retaining the original model parameters to preserve pre-learned knowledge, while adjusting specific additional tensors or layers to cater to new downstream tasks.

Drawing from studies on large language models (LLMs), various Parameter-Efficient Fine-Tuning (PEFT) techniques, such as prefix prompt tuning[25] and LoRA[26], have been proposed to address the issues of catastrophic forgetting and the impractical computational cost associated with fine-tuning. Our goal in this paper is to explore the application of PEFT in scLLMs to enhance their adaptability and effectiveness in the context of cell type identification, a fundamental task in single-cell biology. To achieve this, we initially benchmark current open-sourced finetunable scLLMs, including scBERT, GeneFormer and scGPT to identify the need for adaptation and select the best-performing model, scGPT, for further investigation of PEFT strategies. Subsequently, we design two specific tunable prompts for scLLMs and demonstrate their benefits for cell type identification through comparisons with traditional fine-tuning, prefix prompt tuning, and LoRA in the context of NLP foundation models. To the best of our knowledge, this study represents the first expLoRAtion of PEFT in scLLMs and offers a pathway to leverage scLLMs efficiently and effectively in single-cell biology.

### An overview of current scLLMs

At present, scLLMs conceptualize single-cell expression profiles as a form of biological language. This approach treats the cell expression as a sentence, with each gene describing the cell serving as a word. To extract biological meaning from these “cell sentences,” scLLMs incorporate tokenizer, encoder, and pre-trainer modules similar to those found in LLMs, but they are customized to suit the specific characteristics of single-cell expression profiles.

#### Tokenizer

Similar to LLMs, current scLLMs require tokenization of biological words, which involves converting genes into vectors for subsequent learning. However, the key distinction lies in the fact that the tokenizer of scLLMs needs to combine the gene name and its corresponding expression value. Each scLLM maintains a gene vocabulary to assign a unique integer identifier, *id*(*g*_*j*_), to each gene *g*_*j*_ in an input cell *i*. Consequently, the gene token for cell *i* is represented by a vector, 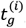, as follows:

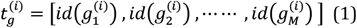

Here, we assume that each input cell comprises *M* genes. Various scLLMs develop their own vocabulary based on their training corpus. In cases where input genes do not align with the predefined vocabulary, scLLMs handle them by utilizing a special padding token to ignore them. Unlike traditional LLMs, scLLMs also need to incorporate each gene’s expression value into gene tokens. A prevalent approach, utilized by scBERT and scGPT, involves categorizing the raw or normalized expression data 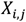 of cell *i* into *k* discrete bins as

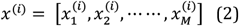

where

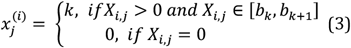

Then, two embedding layers, denoted as *emb*_*g*_ and *emb*_*v*_, are utilized to embed the gene names and gene expression values, respectively, as below

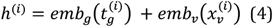

In an exceptional approach, Geneformer ranks genes in descending order according to their expression levels and utilizes positional encoding, as described in reference[27] for LLMs, to embed gene expression values instead.

#### Encoder

The current scLLMs utilize the Transformer architecture to encode gene relationships, drawing on its success in LLMs. This architecture involves stacking *n* transformer blocks[28] (*n* = 6 in scBERT, *n* = 6/12 in Geneformer and *n* = 12 in scGPT), each comprising a self-attention layer, layer normalization, and a Multilayer Perceptron (MLP). This setup is designed to capture interrelated gene patterns, allowing the learned gene embedding 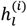 to be computed as follows,

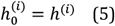

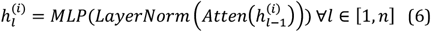

The representation of a cell, treated as a sequence of genes, is generated by pooling all learned gene-level representations *h*^(*i*)^ in scBERT and Geneformer. In the case of scGPT, a special gene-level token < *cls* > is placed as *h*^(*i*+1)^, allowing the models to learn an adaptive gene pooling operation through the self-attention mechanism[28] in transformer blocks.

#### Pretrainer

scLLMs pretrain their models using the Masked Language Model (MLM) objective to encourage the learning of gene contextual features. This objective involves randomly masking certain non-zero gene tokens and predicting the original tokens based on the context provided by the non-masked gene tokens. Typically, a Multilayer Perceptron (MLP) is employed as a decoder to yield the estimated gene tokens. The learning objective can be defined as follows:

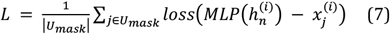

In the equation, *U*_*mask*_ represents the set of masked non-zero genes, and 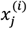 denotes their actual gene expression values. Optional loss functions include Cross Entropy[29], Mean Square Error[29], and Mean Absolute Error[29], among others. The pretraining process involves utilizing multiple single-cell atlases to support these large models. For example, scGPT was trained on 33 million cells across various tissues collected from CELLxGENE[30]; scBERT is grounded in the diverse PanglaoDB[31] with over 1.1 million cells; and Geneformer relies on 29.9 million transcriptomes from the Genecorpus-30M[11].

### Proposed PEFT strategies for scLLMs

#### Gene token prompt

In scLLMs, gene tokens encompass not only gene names but also gene expression values, which can vary across different datasets due to batch effects, leading to out-of-distribution issues. To address this, we have developed a tunable prompt to align the distributions of query and pretrained cell expressions, enabling the projection of all gene embeddings into an optimal format within the tokenizer. The gene token prompt functions as an autoencoder-like adapter layer positioned on top of the gene expression embedding layer. This layer employs a combination of an MLP and a Rectified Linear Unit (RELU) activation to compress d-dimensional gene embeddings into a more compact s-dimensional format (*s* << *d*). Subsequently, another MLP is utilized to recover this into an adaptive d-dimensional gene embedding. As a result, equation (4) is modified as follows:

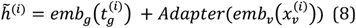

The output gene embedding 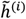 is then fed into the subsequent Transformer-based gene relationship encoder. Throughout training, modifications are applied to this adapter layer while maintaining the native scLLM unchanged. This adapter layer, designed to improve the compatibility of gene tokens, is denoted as the ‘Gene token prompt,’ as illustrated in Figure 1a.

**Figure 1.**
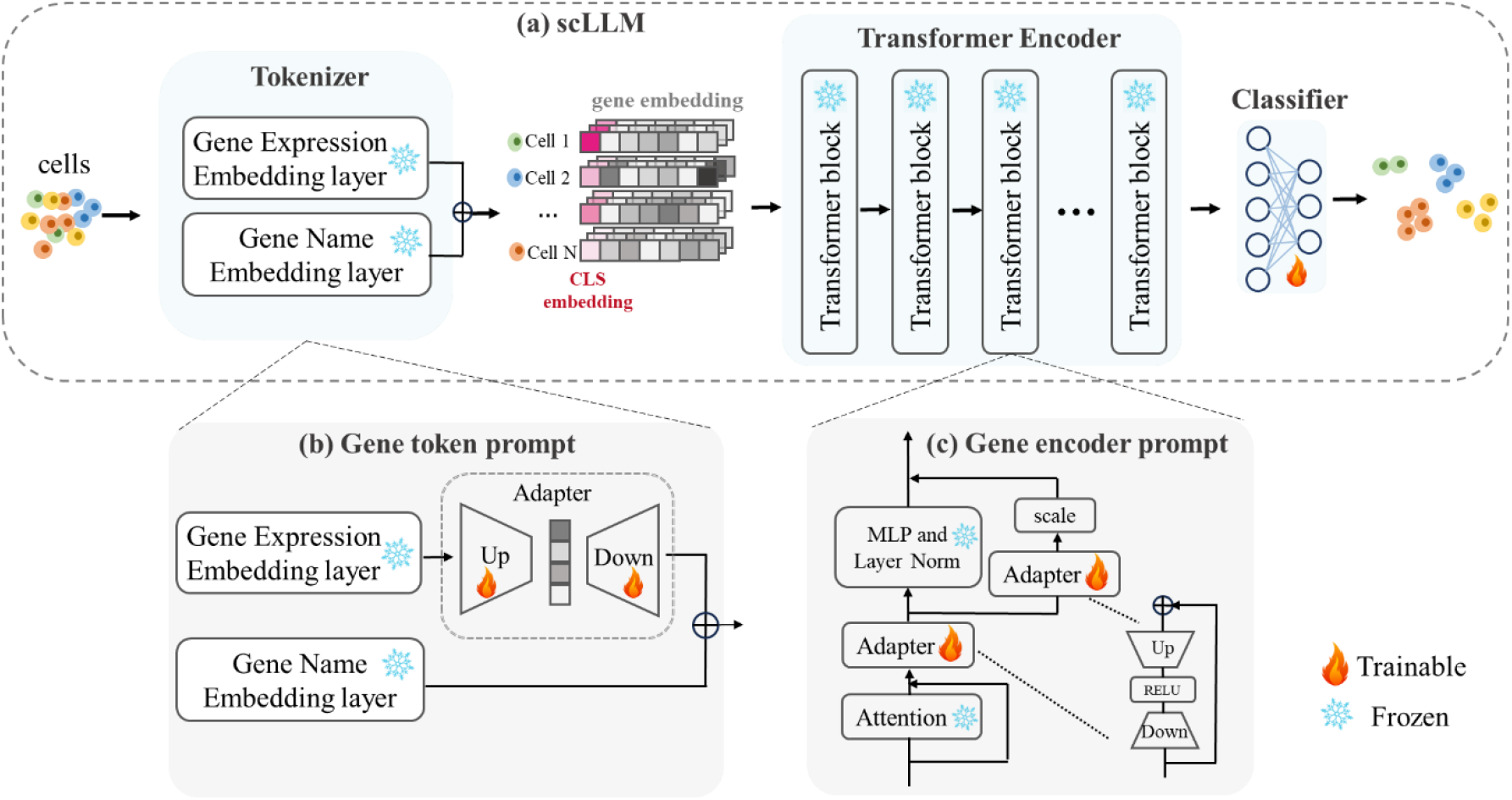
Overview of the two proposed PEFT strategies. (a) A typical scLLM’s architecture covers a tokenizer to encode gene name and gene expression value from a cell to yield gene token embedding, a transformer-based encoder to learn gene relationships across all genes in a cell, and a classifier to decode the gene embedding from encoder to a specific cell type. (b) Gene token prompt: An encoder-decoder configuration adapter that processes the input gene expression profile. During training process, only the adapter undergoes update, while the pretrained scLLM is fixed. (c) Gene encoder prompt: adjustable scale and adapter modules to encoder for adapting gene embedding in gene relationship modeling. Only the parameters of the adapters are updated in training while keeping scGPT parameters frozen.

**Figure 2.**
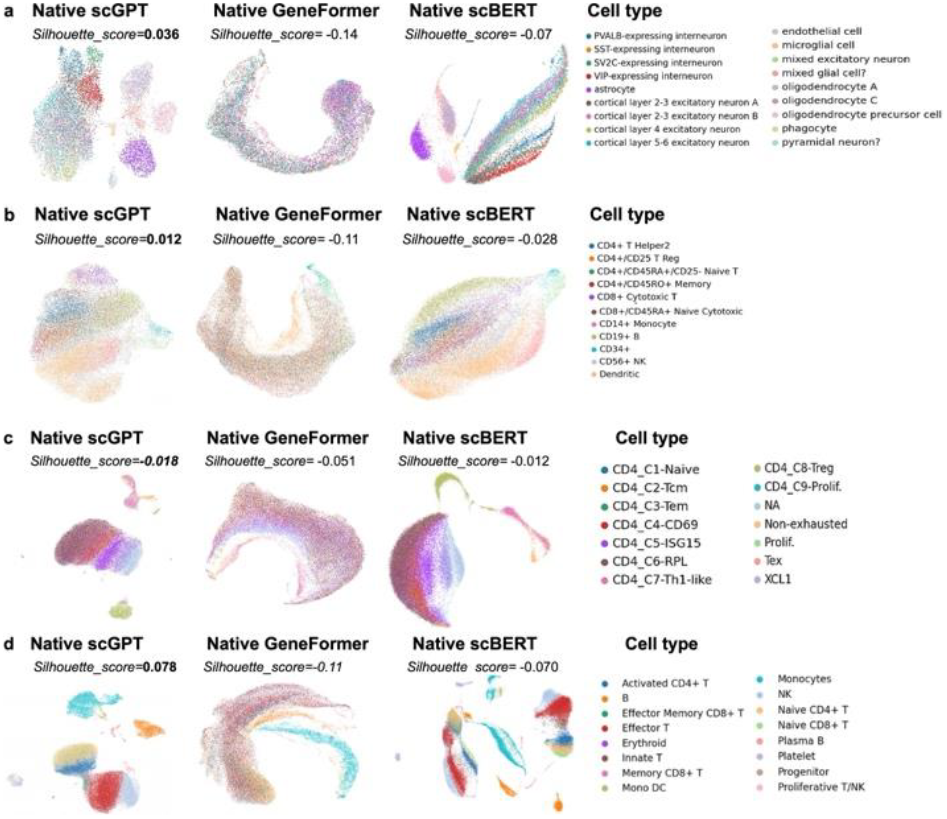
UMAP visualizations of embeddings from scLLMs on (a) M.S., (b) NSCLC, (c) Zheng68k, and (s) COVID-19 dataset. Bold silhouette scores are the best ones across the three scLLMs.

#### Gene encoder prompt

Expanding on a strategy influenced by Wu et al.[32], we incorporated two adapters within the Transformer layers in the targeted scLLM, as depicted in Figure 1b. These adapter layers serve to align the acquired gene relationships from the query data with pretrained universal patterns, thereby inheriting the pretrained knowledge and averting catastrophic forgetting. Additionally, the adapters utilize an autoencoder structure, functioning as a Gene encoder prompt to project the gene embeddings from native Transformers to adaptive subspaces and subsequently reconstruct an optimal gene embedding within those subspaces. Consequently, Equation (6) is revised as:

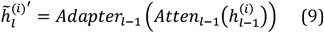

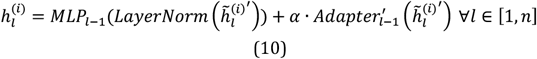

Here, *α* represents a scale factor (in our study, we set *α* = 0.5 by default). We have termed this the “Gene encoder prompt” because it customizes gene context embeddings in the gene encoder section for specific tasks. Throughout the training process, only the adapters will be updated.

#### Finetuning and evaluation settings

When we customized scLLMs by traditional finetuning or proposed PEFT strategies, we employed the Adam optimizer[33], initializing with a learning rate of 10^−5^. As our objective function, we adopted the widely accepted cross entropy loss[29]. As the training process progresses and the loss diminishes, the learning rate is adaptively decreased, minimizing the risk of bypassing the global optimum. We specified the maximum number of training epochs at 100. Among all epochs, the checkpoint exhibiting the minimal loss on the validation set was retained as the fine-tuned model. When incorporating prompt-based learning, all scLLM model parameters were frozen; only the additional tunable tensors were adjusted in response to the gradient of the loss function.

Once the model finished customization, considering the nature of imbalanced cell type distribution, we calculated the metrics including accuracy, precision, recall and weighted F1 score[34] to assess the performance of different approaches. When comparing the power of different finetuning strategies, we further export cell embeddings from tuned scLLMs and compute their Silhouette index to indicate the quality of cell embeddings generated by these strategies.

#### Data preparation

Four datasets were included in our evaluation for fair comparisons as shown in Table 1. These datasets were not involved in the pretraining of current scLLMs. Among them, M.S.[12] and Zheng68k[35] were used to evaluate scGPT and scBERT papers, respectively. We reused them in our evaluation to investigate the reproducibility of the involving scLLMs. Hence, we kept the normalized gene expression profiles, highly variable gene filtering, and data split made by scGPT and scBERT. Also, compared to most native scLLMs pretraining on data from healthy humans, these two datasets cover healthy and diseased conditions suitable for examining the adaptability from different PEFT strategies. Besides, two additional datasets NSCLC[36] and COVID-19[37] from independent studies were employed into our further evaluation. These datasets were all collected from patients with high sequencing genes and diverse cell types, offering clear annotations for our experiments.

**Table 1.** Overview of Datasets involving our evaluation.

| Dataset | Source | Condition | # of genes per cell | # of cells | # of cell types | Data Partition |
| --- | --- | --- | --- | --- | --- | --- |
| M.S. | Brain tissue | Disease and control | 3,000 | 21,312 | 18 | Provided by scGPT[12] |
| Zheng68k | Peripheral blood mononuclear cells | normal | 19,379 | 68,450 | 11 | Provided by scBERT[10] |
| NSCLC | lung cancer cells | disease | 17,022 | 77,030 | 14 | Split by patients |
| COVID-19 | Blood cells | disease | 18,823 | 88,374 | 16 | Split by patients |

**Table 2.** Performance of cell type identification using native scLLMs and popular tools. Bold value represents the highest score among the methods

| Dataset | Method | Accuracy | Precision | Recall | F1-Score |
| --- | --- | --- | --- | --- | --- |
| M.S. | SingleR | 0.580 | 0.776 | 0.663 | 0.615 |
|  | Seurat | **0.686** | **0.820** | **0.812** | **0.805** |
|  | scGPT | 0.595 | 0.777 | 0.728 | 0.734 |
|  | Geneformer | 0.283 | 0.235 | 0.532 | 0.388 |
|  | scBERT | 0.534 | 0.742 | 0.641 | 0.663 |
| Zheng68k | SingleR | 0.543 | 0.571 | 0.377 | 0.398 |
|  | Seurat | **0.553** | **0.674** | **0.662** | **0.645** |
|  | scGPT | 0.525 | 0.632 | 0.639 | 0.625 |
|  | Geneformer | 0.447 | 0.512 | 0.588 | 0.561 |
|  | scBERT | 0.452 | 0.580 | 0.569 | 0.549 |
| NSCLC | SingleR | **0.693** | 0.670 | 0.669 | 0.649 |
|  | Seurat | 0.691 | 0.678 | 0.767 | 0.718 |
|  | scGPT | 0.639 | **0.762** | **0.771** | **0.759** |
|  | Geneformer | 0.545 | 0.701 | 0.645 | 0.679 |
|  | scBERT | 0.583 | 0.736 | 0.735 | 0.718 |
| COVID-19 | SingleR | 0.909 | 0.928 | 0.921 | 0.922 |
|  | Seurat | **0.913** | **0.935** | **0.935** | **0.934** |
|  | scGPT | 0.832 | 0.875 | 0.864 | 0.866 |
|  | Geneformer | 0.684 | 0.769 | 0.732 | 0.752 |
|  | scBERT | 0.735 | 0.855 | 0.839 | 0.837 |

To conduct fair evaluation, we randomly split training and testing sets from these datasets by 8:2 of patient samples and further left one patient out from training set as validation data when training the model. Before training, we applied log1p normalization to their expression values and selected top 2000 highly variable genes as input to keep consistent with the most widely used Seurat protocol[38].

## Results

### Comparison of native scLLMs on cell type identification

In our investigation, we initiated a comparative analysis between scLLMs and two conventional cell type annotation tools, SingleR and Seurat. These conventional tools function by mapping cell types from an annotated reference set to a query set based on raw expression profiles, typically favoring cells with high similarity. In parallel, we applied a similar mapping strategy using generative embeddings from open-sourced scLLMs, including scGPT, Geneformer, and scBERT, to perform cell type identification in a zero-shot setting. For a fair comparison, we designated split training sets as references and testing sets as queries, running all experiments using the default parameters provided by each method.

Among the five methods evaluated, Seurat demonstrated robust performance across three datasets, evidenced by its high precision and recall, signifying its capacity to account for the biological variability present in raw single-cell data. Conversely, SingleR exhibited limitations in precision and recall on datasets M.S, Zheng68k, and NSCLC, suggesting its potential needs of denoising preprocessing steps. Notably, the native representations derived from scLLMs were outperformed by these conventional tools except for scGPT’s performance on the NSCLC dataset. This suggests that the intrinsic representations of scLLMs lack the discerning power for cell type labeling, potentially because these datasets encompass diseased samples, which exhibit a distributional shift compared to the pretraining data derived from normal conditions. Furthermore, their pretraining objectives were not explicitly oriented towards cell type identification. Consequently, these models’ representations are not inherently task-specific, necessitating further adaptation for downstream tasks, a finding that is corroborated by existing literature[17-18].

Among the scLLMs, scGPT showed enhanced robustness across the four datasets compared to scBERT and GeneFormer, which may be attributed to its larger model size and extensive pretraining data. This observation aligns with that larger architectures generally possess greater capacity. However, the escalation in model size imposes a significant computational burden, particularly for large-scale single-cell analyses. These constraints motevated our pursuit of developing efficient adaptation approaches that could facilitate scLLMs in single-cell biology. With scGPT emerging as the most capable model among the three scLLMs, we selected it to demonstrate the efficacy of our proposed PEFT strategies in the following experiments.

To further evaluate the representational capabilities of the three scLLMs, we employed Uniform Manifold Approximation and Projection (UMAP) to translate their high-dimensional embeddings into a two-dimensional space, subsequently color-coding the resulting plots by cell type. We also computed silhouette scores as a quantitative measure of the quality of the scLLM representations, where values range from −1 to 1 with higher scores indicating more distinct clustering of cell types.

The UMAP projections and corresponding silhouette scores revealed that scGPT produced varied cluster densities and degrees of separation. In certain instances, distinct cell types were more delineated, suggesting a comparatively superior representational capability. While scGPT’s silhouette scores were relatively higher, indicating some promise, they did not reach a level of satisfaction. The UMAP plots for scBERT demonstrated a degree of distinctiveness, despite of less pronounced than that of scGPT, as reflected by its lower silhouette scores. Conversely, Geneformer’s UMAP visualizations and silhouette scores did not exhibit a discernible clustering pattern among different cell types, consistent with its performance metrics. These UMAP analyses collectively highlight the existing limitations of the three scLLMs in zero-shot cell type annotation, underscoring the need for further refinement to enhance their discriminative capacity in this application.

### Comparison of proposed PEFT strategies and other finetuning approaches

We then integrated our innovative Parameter-Efficient Fine-Tuning (PEFT) strategies to fine-tune scGPT using identical training data, subsequently evaluating the tuned models on the test set. For benchmarking purposes, we juxtaposed our PEFT strategies against the conventional fine-tuning approach, which encompasses a comprehensive update of the encoder and classifier, as well as an alternative approach that fine-tunes the classifier only, leaving the encoder unchanged. While the former represents the standard yet resource-intensive method in deep learning, its feasibility diminishes with increasing model sizes. The latter, conversely, stands as the least computationally demanding refinement technique. In addition to these, we examined our PEFT strategies alongside two prevalent prompt-based learning techniques: prefix prompting and LoRa. We summarized the comparative performance across all cell types in Table 3, with detailed results for individual cell types provided in Supplementary Tables 13-36. It is important to note that we explored multiple hyperparameter configurations for our PEFT strategies and competing approaches, with a comprehensive account of these variations detailed in Supplementary Tables 37-39. The results reported in Table 3 reflect the outcomes from the optimally performing hyperparameter sets.

**Table 3.** Performance of scGPT with proposed PEFT strategies and other adaptation approaches. Italic values denote the best performance from Table 2 as baseline. Bold values represent the best metric values across all involved approaches.

| Dataset | Method | Trainable Parameters | Accuracy | Precision | Recall | F1-Score |
| --- | --- | --- | --- | --- | --- | --- |
| M.S. | *Seurat* | N/A | 0.686 | 0.820 | 0.812 | 0.805 |
|  | Native scGPT | N/A | 0.595 | 0.777 | 0.728 | 0.734 |
|  | Full finetune | 51M | 0.708 | 0.857 | 0.848 | 0.845 |
|  | Finetune classifier | 0.53M | 0.650 | 0.806 | 0.801 | 0.797 |
|  | Prefix prompt | 0.92M | 0.743 | 0.815 | 0.797 | 0.802 |
|  | LoRA prompt | 1.27M | 0.747 | 0.830 | 0.817 | 0.821 |
|  | Gene token prompt | 0.66M | 0.790 | 0.875 | 0.842 | 0.852 |
|  | Gene encoder prompt | 2.11M | **0.795** | **0.884** | **0.870** | **0.874** |
| Zheng68k | *Seurat* | N/A | 0.553 | 0.674 | 0.662 | 0.645 |
|  | Native scGPT | N/A | 0.525 | 0.632 | 0.639 | 0.625 |
|  | Full finetune | 51M | 0.681 | 0.771 | 0.654 | 0.679 |
|  | Finetune classifier | 0.53M | 0.546 | 0.681 | 0.677 | 0.656 |
|  | Prefix prompt | 0.92M | 0.590 | 0.735 | 0.484 | 0.528 |
|  | LoRA prompt | 1.27M | 0.582 | 0.730 | 0.499 | 0.545 |
|  | Gene token prompt | 0.66M | 0.678 | 0.791 | 0.682 | 0.708 |
|  | Gene encoder prompt | 2.11M | **0.696** | **0.809** | **0.811** | **0.807** |
| NSCLC | *Seurat* | N/A | 0.691 | 0.678 | 0.767 | 0.718 |
|  | Native scGPT | N/A | 0.639 | 0.762 | 0.771 | 0.759 |
|  | Full finetune | 51M | 0.772 | 0.873 | 0.872 | 0.873 |
|  | Finetune classifier | 0.53M | 0.712 | 0.813 | 0.816 | 0.812 |
|  | Prefix prompt | 0.92M | 0.773 | 0.802 | 0.759 | 0.769 |
|  | LoRA prompt | 1.27M | 0.746 | 0.809 | 0.779 | 0.786 |
|  | Gene token prompt | 0.66M | **0.886** | 0.878 | 0.869 | 0.871 |
|  | Gene encoder prompt | 2.11M | 0.883 | **0.884** | **0.874** | **0.876** |
| COVID-19 | *Seurat* | N/A | 0.913 | 0.935 | 0.935 | 0.934 |
|  | Native scGPT | N/A | 0.832 | 0.875 | 0.864 | 0.866 |
|  | Full finetune | 51M | 0.921 | 0.939 | 0.935 | 0.936 |
|  | Finetune classifier | 0.53M | 0.833 | 0.915 | 0.916 | 0.915 |
|  | Prefix prompt | 0.92M | 0.914 | 0.916 | 0.906 | 0.909 |
|  | LoRA prompt | 1.27M | 0.911 | 0.912 | 0.906 | 0.908 |
|  | Gene token prompt | 0.66M | 0.950 | 0.947 | 0.944 | 0.945 |
|  | Gene encoder prompt | 2.11M | **0.957** | **0.949** | **0.946** | **0.947** |

**Table 4.** Comparison of scGPT with different adaptation approaches on the rare cell type from the M.S. dataset. Bold values represent the best metric values across all involved approaches.

| Cell type | Support | Method | Accuracy | Precision | Recall | F1-score |
| --- | --- | --- | --- | --- | --- | --- |
| cortical layer<br>2-3 excitatory<br>neuron A | 314 | SingleR | 0.0489 | 0.3000 | 0.0382 | 0.0678 |
|  |  | Seurat | 0.5809 | 0.3014 | **0.8408** | 0.4437 |
|  |  | Native scGPT | 0.3300 | 0.1345 | 0.6401 | 0.2223 |
|  |  | Full Finetune | 0.4841 | 0.3101 | 0.5955 | 0.4079 |
|  |  | Prefix prompt | 0.3165 | 0.2408 | 0.3535 | 0.2865 |
|  |  | LoRA prompt | 0.4173 | 0.3500 | 0.4459 | 0.3922 |
|  |  | Gene token prompt | 0.5491 | 0.3052 | 0.7484 | 0.4336 |
|  |  | Gene encoder prompt | **0.5620** | **0.3739** | 0.6752 | **0.4813** |
| phagocyte | 82 | SingleR | 0.0478 | 0.6000 | 0.0366 | 0.0690 |
|  |  | Seurat | 0.0160 | 0.2500 | 0.0122 | 0.0233 |
|  |  | Native scGPT | 0.0000 | 0.0000 | 0.0000 | 0.0000 |
|  |  | Full Finetune | 0.0000 | 0.0000 | 0.0000 | 0.0000 |
|  |  | Prefix prompt | 0.4737 | 0.6207 | 0.4390 | 0.5143 |
|  |  | LoRA prompt | 0.5195 | 0.6452 | **0.4878** | 0.5556 |
|  |  | Gene token prompt | **0.5217** | 0.7358 | 0.4756 | **0.5778** |
|  |  | Gene encoder prompt | 0.3986 | **0.8000** | 0.3415 | 0.4786 |

**Table 5.**
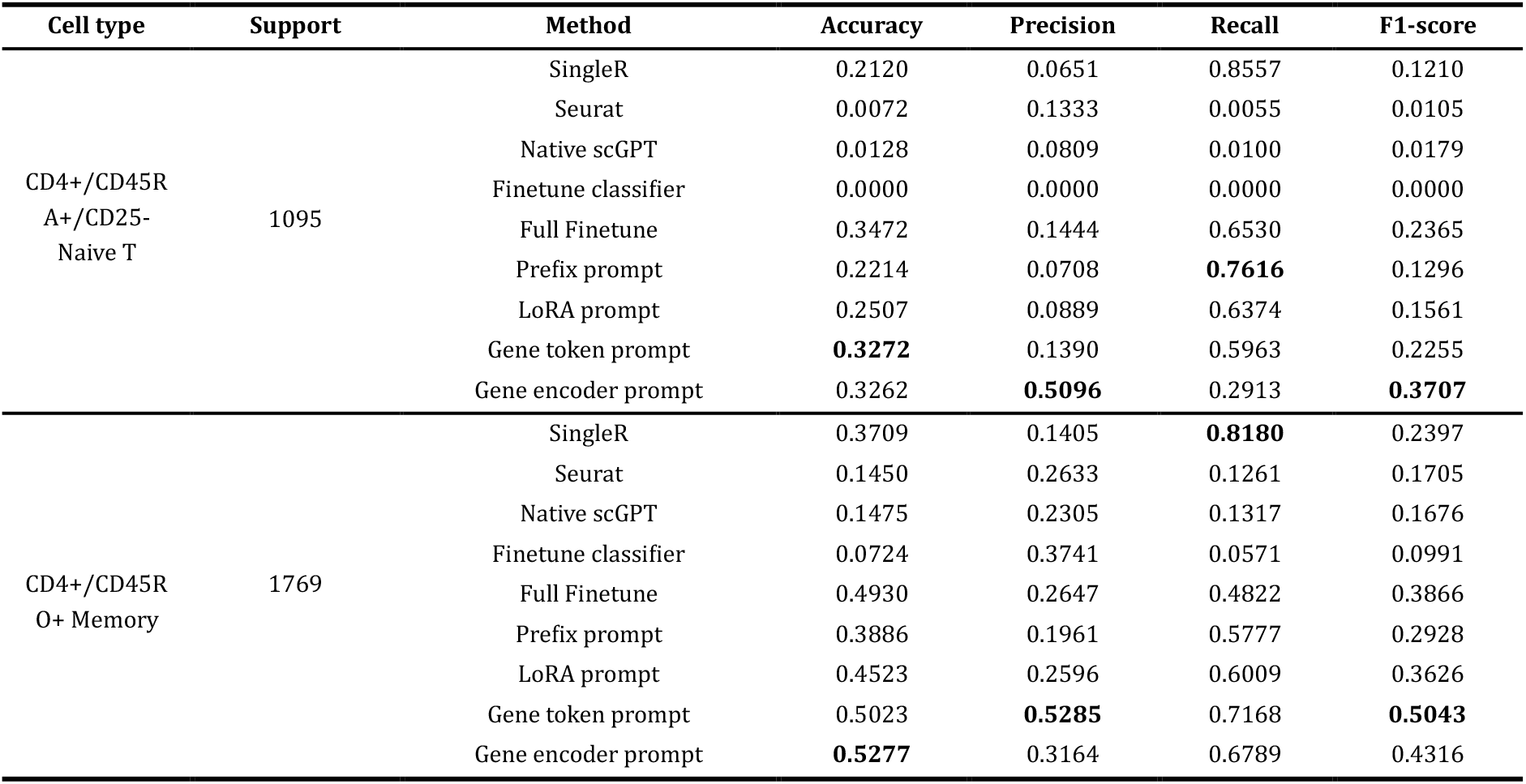
Comparison of scGPT with different adaptation approaches on the ambiguous cell type from the Zheng68k dataset. Bold values represent the best metric values across all involved approaches.

The results display a marked enhancement in model performance post-fine-tuning compared to their native counterparts, underlining the critical role of model adaptation in cell type identification using scGPT. However, full fine-tuning, despite its effectiveness, is computationally exhaustive, necessitating updates across all transformer blocks, encompassing over 50 million parameters—a scale that is impractible for routine single-cell analysis. The classifier-only fine-tuning approach, while resource-conservative, fails to adequately adapt the scLLMs as evidenced by its relatively poorer performance metrics.

Our PEFT strategies, by fine-tuning a subset of newly added parameters, adeptly capture task-specific information within scGPT, leading to consistently superior performance across a spectrum of metrics, including Accuracy, Precision, Recall, and F1-score. These strategies operate at the tokenizer and encoder levels, tuning raw input gene expression values and the learned gene relationships from the Transformer blocks to fit the task-specific demands. Consequently, not only do they enhance performance relative to Seurat and native scGPT, but they also outstrip the full fine-tuning approach while saving up to 90% trainable parameters. Between our two PEFT strategies, the Gene token prompt slightly trails the Gene encoder prompt, yet it offers a further reduction of trainable parameters by one-third. Despite their popularity in NLP models, the prefix prompt and LoRa prompt did not fare as well in our experiments. The prefix prompt, which appends pseudo tokens to the input, potentially disrupts the biological signal within the gene expression data. LoRa’s requirement for a sufficiently large rank number to capture complex biological patterns also appeared to be a limiting factor in our context.

Figure 3 presents the UMAP visualizations derived from the various adaptation approaches, showcasing the enhanced delineation of cell representations achieved by the fine-tuned scGPT. It is apparent from the visualizations that all adaptation methods yield representations that are superior to those generated by the native scGPT model. Notably, our PEFT strategies outperform the alternatives, producing more cohesive clustering as evidenced by UMAP visualizations and higher Silhouette scores. This underscores the efficacy of the PEFT-induced prompts in fostering a refined predictive capability for cell type identification. In contrast, the prefix prompt approach appears to falter in generating effective cell representations. Meanwhile, LoRA prompts, though not as effective as PEFT strategies, still deliver relatively promising representation quality.

**Figure 3.**
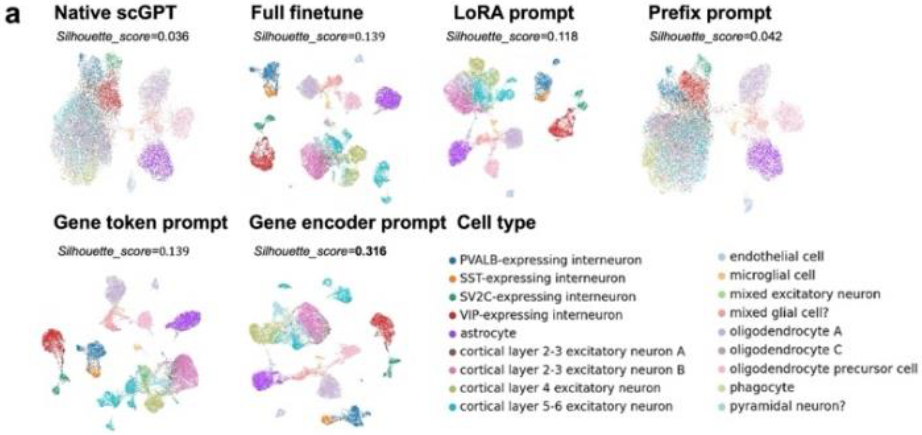
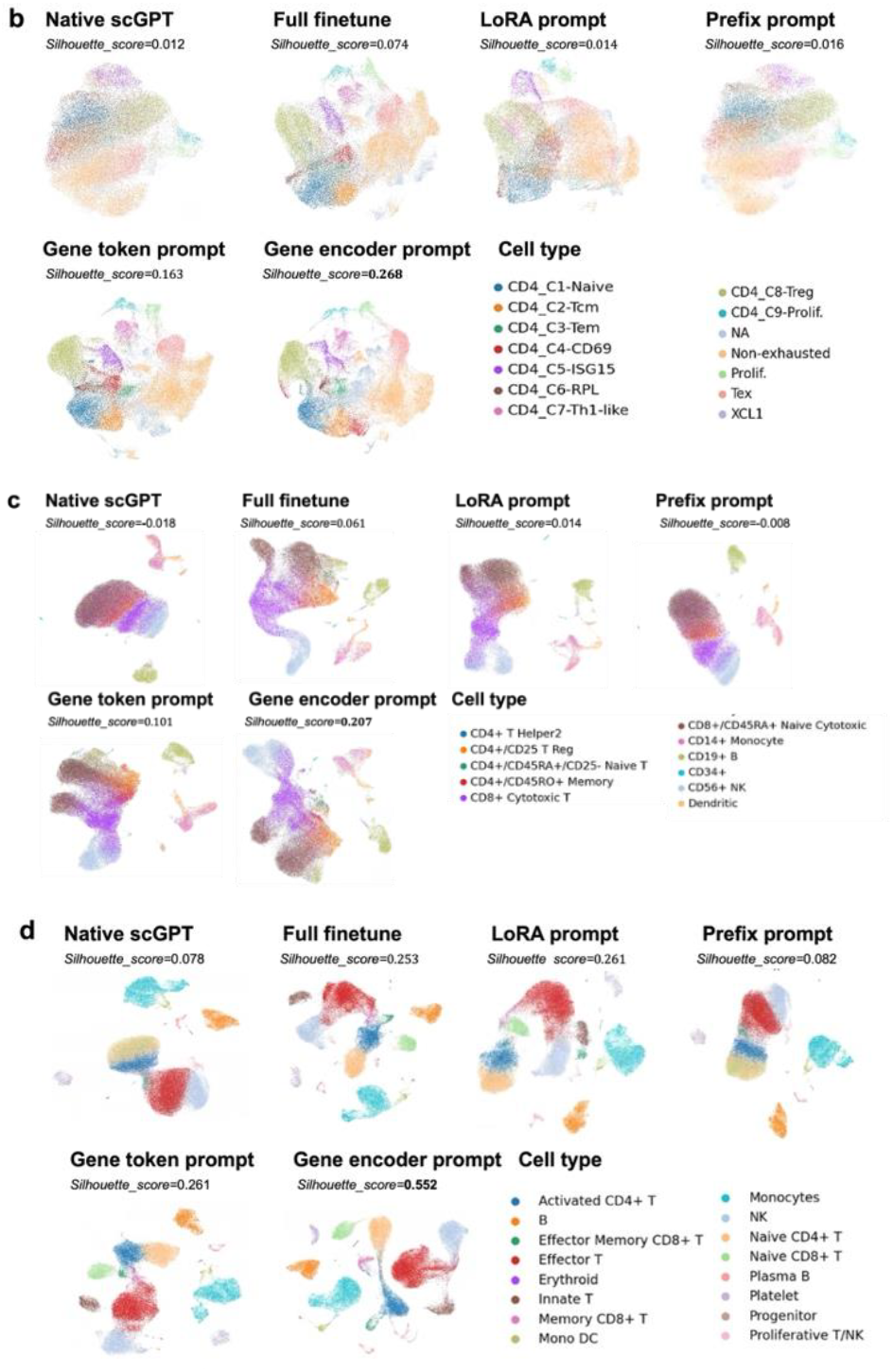
UMAP visualizations of scGPT with different adaptation approaches on (A) M.S., (B) NSCLC, (C) Zheng68k, and (D) COVID-19 Datasets.

### Performance of proposed PEFT strategies and other finetuning approaches on challenging cell types

In cell type identification task, addressing “hard cases” – cell types that are challenging to classify due to rarity and overshadowing by dominant populations – is vital for understanding cellular diversity, especially in complex diseases like cancer. These hard cases, exemplified by Naive T (CD4+/CD45RA+/CD25-) and Memory T (CD4+/CD45RO+) cells in the Zheng68k dataset, pose challenges as traditional feature extraction often fails to capture the subtle differences between closely related subtypes. This challenge is primarily attributed to the potential inadequacy of features extracted from the data, which may not sufficiently represent the nuanced differences between these T cell subtypes.

Our study conducts a comprehensive comparative analysis on datasets such as M.S., focusing on these elusive cell types. The results, highlighted in Table 6, reveal the significant efficacy of the Gene encoder prompt in improving classification. This PEFT approach effectively tailors the model to capture specific data traits, such as unique gene expression patterns, enhancing the model’s precision in identifying hard-to-classify cell types and underscoring the importance of methodological innovation in single-cell genomics.

**Table 6.** Comparison of scGPT with different adaptation approaches on the rare cell type from the NSCLC dataset. Bold values represent the best metric values across all involved approaches.

| Cell type | Support | Method | Accuracy | Precision | Recall | F1-score |
| --- | --- | --- | --- | --- | --- | --- |
| XCL1 | 52 | SingleR | 0.0768 | 0.0240 | 0.2885 | 0.0442 |
|  |  | Seurat | 0.0250 | 0.2500 | 0.0192 | 0.035 |
|  |  | Native scGPT | 0.0000 | 0.0000 | 0.0000 | 0.0000 |
|  |  | Finetune classifier | 0.0000 | 0.0000 | 0.0000 | 0.0000 |
|  |  | Full Finetune | 0.0000 | 0.0000 | 0.0000 | 0.0000 |
|  |  | prefix prompt | 0.0987 | 0.0302 | 0.4038 | 0.0561 |
|  |  | LoRA prompt | 0.3357 | 0.1341 | 0.6731 | 0.2236 |
|  |  | Gene token prompt | **0.5926** | 0.3121 | **0.8462** | 0.4560 |
|  |  | Gene encoder prompt | 0.5839 | **0.3390** | 0.7692 | **0.4706** |

## Conclusion and Discussion

In this study, we proposed two PEFT strategies for enhancing the adaptability of scLLMs. Our approach introduces a novel concept of tunable Gene token and Gene encoder prompts within scLLMs, while maintaining the integrity of the original model parameters during their adaptation to specific downstream tasks. Through rigorous evaluation using scGPT across four benchmark datasets for cell type identification, our strategies have demonstrated improvements in accuracy, precision, recall, and F1 score, achieved with substantially lower computational demands. Notably, our PEFT strategies have shown marked enhancements in identifying rare cell types and complex cell types on these datasets, thereby revealing the power of scLLMs in such critical scenarios.

As an initial investigation into the benefits of PEFT approaches applied to scLLMs for single-cell analysis, this work has exclusively utilized scGPT as the representative scLLM. Further research is warranted to establish a comprehensive benchmark across additional open-sourced scLLMs, such as scBERT and Genefomer, to assess the robustness of these strategies. Moreover, future endeavors will aim to expand the application of PEFT techniques beyond cell type identification to other fundamental single-cell tasks, including batch correction, perturbation response analysis, and gene marker detection, thus enhancing the utility of scLLMs in the single-cell community.

Additionally, the development of a dedicated PEFT toolkit for emerging scLLMs stands as an objective to facilitate and streamline research endeavors involving scLLMs and PEFT strategies in single-cell biology. Exploring the combination of various PEFT strategies may yield a comprehensive solution for scLLMs, particularly for complex tasks such as network inference. Inspired by the ‘chain of thoughts’ methodology in LLMs, these intricate tasks could be deconstructed into sequential subtasks—ranging from cell type to network identification—and tackled using targeted prompts to effectively harness the power of scLLMs.

Lastly, the pursuit of integrating scLLMs with large foundational models from disparate modalities, such as imaging or proteomics, is expected. The aim is to cultivate a more holistic understanding of cellular behaviors and interactions at a multimodal scale, which remains a promising yet challenging frontier in the field.

## Data and code availability

All data used in this study are publicly available. The published Zheng68k dataset was downloaded at https://support.10xgenomics.com/single-cell-gene-expression/datasets(SRP073767)[35]. The published M.S. dataset s were downloaded from Github at https://github.com/bowang-lab/scGPT/tree/main/data/. The NSCLC dataset was downloaded from https://www.ncbi.nlm.nih.gov/geo/query/acc.cgi?acc=GSE179994. The COVID-19 dataset was downloaded https://doi.org/10.6084/m9.figshare.16922467.v1. The source code is freely available on Github (https://github.com/laolintou/scPEFT.git)

## Acknowledgements

We would like to thank Mengran Zhao and Baichuan Jin for their technical support. We also would like to acknowledge the valuable contribution from OpenAI’s GPT-4 model for aiding in language editing. This work was funded by the National Institutes of Health [R35-GM126985].

## Appendix

There is one additional file containing Supplementary Tables 1-39.

## Notes

### Competing Interest Statement

The authors have declared no competing interest.

