## Supplementary Tables 1-39 for "Parameter-Efficient Fine-Tuning Enhances Adaptation of Single Cell Large Language Model for Cell Type Identification"

**Supplementary Table 1. Performance of SingleR across cell types on the MS dataset**

| Cell type | Precision | Recall | F1-score | Support |
|---|---|---|---|---|
| PVALB-expressing interneuron | 0.9189 | 0.9032 | 0.9110 | 878 |
| PVALB-expressing interneuron | 0.8617 | 0.6279 | 0.7265 | 258 |
| SV2C-expressing interneuron | 0.7109 | 0.9651 | 0.8187 | 344 |
| VIP-expressing interneuron | 0.9965 | 0.7288 | 0.8419 | 1158 |
| astrocyte | 0.8956 | 0.9872 | 0.9392 | 1330 |
| cortical layer 2-3 excitatory neuron A | 0.3000 | 0.0382 | 0.0678 | 314 |
| cortical layer 2-3 excitatory neuron B | 0.5267 | 0.7369 | 0.6144 | 2178 |
| cortical layer 4 excitatory neuron | 0.8217 | 0.1961 | 0.3167 | 1504 |
| cortical layer 5-6 excitatory neuron | 0.9316 | 0.1259 | 0.2218 | 1732 |
| endothelial cell | 0.9655 | 0.9882 | 0.9767 | 170 |
| microglial cell | 0.8699 | 0.9068 | 0.8880 | 118 |
| mixed excitatory neuron | 0.4000 | 0.0324 | 0.0600 | 185 |
| mixed glial cell | 0.8525 | 0.3630 | 0.5092 | 573 |
| oligodendrocyte A | 0.7627 | 0.9967 | 0.8642 | 1216 |
| oligodendrocyte C | 0.0000 | 0.0000 | 0.0000 | 34 |
| oligodendrocyte precursor cell | 0.9920 | 0.8580 | 0.9201 | 866 |
| phagocyte | 0.6000 | 0.0366 | 0.0690 | 82 |
| pyramidal neuron | 0.1666 | 0.9640 | 0.2840 | 528 |

**Supplementary Table 2. Performance of SingleR across cell types on the Zheng68k dataset**

| Cell type | Precision | Recall | F1-score | Support |
|---|---|---|---|---|
| CD4+ T Helper2 | 0.0552 | 0.1311 | 0.0777 | 61 |
| CD4+/CD25 T Reg | 0.6399 | 0.1669 | 0.2648 | 3672 |
| CD4+/CD45RA+/CD25- Naive T | 0.0651 | 0.8557 | 0.1210 | 1095 |
| CD4+/CD45RO+ Memory | 0.1405 | 0.8180 | 0.2397 | 1769 |
| CD8+ Cytotoxic T | 0.9019 | 0.1103 | 0.1965 | 12340 |
| CD8+/CD45RA+ Naive Cytotoxic | 0.6719 | 0.1681 | 0.2689 | 10161 |
| CD14+ Monocyte | 0.8128 | 0.7981 | 0.8054 | 1714 |
| CD19+ B | 0.9799 | 0.6354 | 0.7709 | 3524 |
| CD34+ | 0.9709 | 0.6993 | 0.8130 | 143 |
| CD56+ NK | 0.8159 | 0.9106 | 0.8606 | 5290 |
| Dendritic | 0.7098 | 0.6826 | 0.6959 | 1301 |

**Supplementary Table 3. Performance of SingleR across cell types on the NSCLC dataset**

| Cell type | Precision | Recall | F1-score | Support |
|---|---|---|---|---|
| CD4_C1-Naive | 0.8055 | 0.6253 | 0.7040 | 3728 |
| CD4_C2-Tcm | 0.3363 | 0.8905 | 0.4882 | 1982 |
| CD4_C3-Tem | 0.3733 | 0.7385 | 0.4959 | 1205 |
| CD4_C4-CD69 | 0.4069 | 0.7771 | 0.5342 | 1539 |
| CD4_C5-ISG15 | 0.6214 | 0.8031 | 0.7007 | 1402 |
| CD4_C6-RPL | 0.6235 | 0.6712 | 0.6465 | 1034 |
| CD4_C7-Th1-like | 0.6687 | 0.7494 | 0.7068 | 1325 |
| CD4_C8-Treg | 0.8895 | 0.8617 | 0.8754 | 5714 |
| CD4_C9-Prolif. | 0.9372 | 0.8326 | 0.8818 | 484 |
| NA | 0.0000 | 0.0000 | 0.0000 | 4652 |
| Non-exhausted | 0.9123 | 0.6829 | 0.7811 | 11714 |
| Prolif. | 0.8545 | 0.8265 | 0.8403 | 732 |
| Tex | 0.6065 | 0.9540 | 0.7415 | 3132 |
| XCL1 | 0.0240 | 0.2885 | 0.0442 | 52 |

**Supplementary Table 4. Performance of SingleR across cell types on the COVID dataset**

| Cell type | Precision | Recall | F1-score | Support |
|---|---|---|---|---|
| Activated CD4+ T | 0.8115 | 0.9245 | 0.8643 | 5075 |
| B | 0.9997 | 0.9997 | 0.9997 | 2955 |
| Effector Memory CD8+ T | 0.6098 | 0.9744 | 0.7502 | 547 |
| Effector T | 0.9750 | 0.8955 | 0.9336 | 13102 |
| Erythroid | 1.0000 | 0.8060 | 0.8926 | 67 |
| Innate T | 0.7888 | 0.9602 | 0.8661 | 1610 |
| Memory CD8+ T | 0.6221 | 0.6572 | 0.6392 | 849 |
| Mono DC | 0.9730 | 0.9059 | 0.9382 | 478 |
| Monocytes | 0.9919 | 0.9990 | 0.9954 | 5728 |
| NK | 0.8775 | 0.9506 | 0.9126 | 6214 |
| Naive CD4+ T | 0.9695 | 0.8252 | 0.8916 | 5705 |
| Naive CD8+ T | 0.9518 | 0.9713 | 0.9615 | 2338 |
| Plasma B | 0.9880 | 0.9970 | 0.9925 | 331 |
| Platelet | 1.0000 | 1.0000 | 1.0000 | 919 |
| Progenitor | 1.0000 | 0.8917 | 0.9428 | 157 |
| Proliferative T/NK | 0.9254 | 0.8017 | 0.8591 | 464 |

**Supplementary Table 5. Performance of Seurat across cell types on the MS dataset**

| Cell type | Precision | Recall | F1-score | Support |
|---|---|---|---|---|
| PVALB-expressing interneuron | 0.9446 | 0.8941 | 0.9187 | 878 |
| PVALB-expressing interneuron | 0.7533 | 0.8760 | 0.8100 | 258 |
| SV2C-expressing interneuron | 0.9030 | 0.9477 | 0.9248 | 344 |
| VIP-expressing interneuron | 0.9772 | 0.9611 | 0.9691 | 1158 |
| astrocyte | 0.9435 | 0.9910 | 0.9666 | 1330 |
| cortical layer 2-3 excitatory neuron A | 0.3014 | 0.8408 | 0.4437 | 314 |
| cortical layer 2-3 excitatory neuron B | 0.7828 | 0.7663 | 0.7745 | 2178 |
| cortical layer 4 excitatory neuron | 0.7348 | 0.8218 | 0.7759 | 1504 |
| cortical layer 5-6 excitatory neuron | 0.7958 | 0.7200 | 0.7560 | 1732 |
| endothelial cell | 0.9494 | 0.9941 | 0.9713 | 170 |
| microglial cell | 0.9091 | 0.6780 | 0.7767 | 118 |
| mixed excitatory neuron | 0.0000 | 0.0000 | 0.0000 | 185 |
| mixed glial cell | 0.7854 | 0.6387 | 0.7045 | 573 |
| oligodendrocyte A | 0.8042 | 0.9391 | 0.8665 | 1216 |
| oligodendrocyte C | 0.0000 | 0.0000 | 0.0000 | 34 |
| oligodendrocyte precursor cell | 0.9764 | 0.9550 | 0.9656 | 866 |
| phagocyte | 0.2500 | 0.0122 | 0.0233 | 82 |
| pyramidal neuron | 0.9222 | 0.3144 | 0.4689 | 528 |

**Supplementary Table 6. Performance of Seurat across cell types on the Zheng68k dataset**

| Cell type | Precision | Recall | F1-score | Support |
|---|---|---|---|---|
| CD4+ T Helper2 | 0.7503 | 0.9714 | 0.8467 | 1714 |
| CD4+/CD25 T Reg | 0.9397 | 0.6674 | 0.7805 | 3524 |
| CD4+/CD45RA+/CD25- Naive T | 0.9353 | 0.9091 | 0.9220 | 143 |
| CD4+/CD45RO+ Memory | 0.0000 | 0.0000 | 0.0000 | 61 |
| CD8+ Cytotoxic T | 0.4496 | 0.5370 | 0.4895 | 3672 |
| CD8+/CD45RA+ Naive Cytotoxic | 0.1333 | 0.0055 | 0.0105 | 1095 |
| CD14+ Monocyte | 0.2633 | 0.1261 | 0.1705 | 1769 |
| CD19+ B | 0.8577 | 0.9233 | 0.8893 | 5290 |
| CD34+ | 0.7622 | 0.5139 | 0.6139 | 12340 |
| CD56+ NK | 0.5534 | 0.8761 | 0.6783 | 10161 |
| Dendritic | 0.8673 | 0.5527 | 0.6751 | 1301 |

**Supplementary Table 7. Performance of Seurat across cell types on the NSCLC dataset**

| Cell type | Precision | Recall | F1-score | Support |
|---|---|---|---|---|
| CD4_C1-Naive | 0.7471 | 0.8114 | 0.7779 | 3728 |
| CD4_C2-Tcm | 0.4983 | 0.7497 | 0.5987 | 1982 |
| CD4_C3-Tem | 0.6131 | 0.6107 | 0.6119 | 1205 |
| CD4_C4-CD69 | 0.6033 | 0.6186 | 0.6108 | 1539 |
| CD4_C5-ISG15 | 0.8295 | 0.8331 | 0.8313 | 1402 |
| CD4_C6-RPL | 0.7390 | 0.6954 | 0.7165 | 1034 |
| CD4_C7-Th1-like | 0.6544 | 0.8445 | 0.7374 | 1325 |
| CD4_C8-Treg | 0.8077 | 0.9268 | 0.8632 | 5714 |
| CD4_C9-Prolif. | 0.9423 | 0.8430 | 0.8899 | 484 |
| NA | 0.0000 | 0.0000 | 0.0000 | 4652 |
| Non-exhausted | 0.8215 | 0.9666 | 0.8881 | 11714 |
| Prolif. | 0.9003 | 0.8757 | 0.8878 | 732 |
| Tex | 0.8396 | 0.8857 | 0.8620 | 3132 |
| XCL1 | 0.2500 | 0.0192 | 0.0357 | 52 |

**Supplementary Table 8. Performance of Seurat across cell types on the COVID dataset**

| Cell type | Precision | Recall | F1-score | Support |
|---|---|---|---|---|
| Activated CD4+ T | 0.9325 | 0.9017 | 0.9169 | 5057 |
| B | 0.9993 | 0.9997 | 0.9995 | 2955 |
| Effector Memory CD8+ T | 0.8629 | 0.8629 | 0.8629 | 547 |
| Effector T | 0.9468 | 0.9309 | 0.9388 | 13102 |
| Erythroid | 1.0000 | 0.8657 | 0.9280 | 67 |
| Innate T | 0.8190 | 0.9640 | 0.8856 | 1610 |
| Memory CD8+ T | 0.7066 | 0.5701 | 0.6310 | 849 |
| Mono DC | 0.9375 | 0.9414 | 0.9395 | 478 |
| Monocytes | 0.9962 | 0.9944 | 0.9953 | 5728 |
| NK | 0.9046 | 0.8943 | 0.8994 | 6214 |
| Naive CD4+ T | 0.9195 | 0.9607 | 0.9397 | 5705 |
| Naive CD8+ T | 0.9141 | 0.9607 | 0.9368 | 2338 |
| Plasma B | 0.9706 | 0.9970 | 0.9836 | 331 |
| Platelet | 1.0000 | 1.0000 | 1.0000 | 919 |
| Progenitor | 0.9463 | 0.8981 | 0.9216 | 157 |
| Proliferative T/NK | 0.9140 | 0.8707 | 0.8918 | 464 |

**Supplementary Table 9. Performance of native scGPT across cell types on the MS dataset**

| Cell type | Precision | Recall | F1-score | Support |
|---|---|---|---|---|
| PVALB-expressing interneuron | 0.8825 | 0.8212 | 0.8507 | 878 |
| PVALB-expressing interneuron | 0.7109 | 0.5814 | 0.6397 | 258 |
| SV2C-expressing interneuron | 0.8842 | 0.9099 | 0.8968 | 344 |
| VIP-expressing interneuron | 0.9641 | 0.8584 | 0.9082 | 1158 |
| astrocyte | 0.9365 | 0.9429 | 0.9397 | 1330 |
| cortical layer 2-3 excitatory neuron A | 0.1345 | 0.6401 | 0.2223 | 314 |
| cortical layer 2-3 excitatory neuron B | 0.6392 | 0.7158 | 0.6753 | 2178 |
| cortical layer 4 excitatory neuron | 0.7378 | 0.6137 | 0.6701 | 1504 |
| cortical layer 5-6 excitatory neuron | 0.7506 | 0.5110 | 0.6080 | 1732 |
| endothelial cell | 0.8852 | 0.9529 | 0.9178 | 170 |
| microglial cell | 1.0000 | 0.1186 | 0.2121 | 118 |
| mixed excitatory neuron | 0.2308 | 0.0324 | 0.0569 | 185 |
| mixed glial cell | 0.5878 | 0.7243 | 0.6489 | 573 |
| oligodendrocyte A | 0.9268 | 0.9161 | 0.9214 | 1216 |
| oligodendrocyte C | 0.0000 | 0.0000 | 0.0000 | 34 |
| oligodendrocyte precursor cell | 0.9756 | 0.8303 | 0.8971 | 866 |
| phagocyte | 0.0000 | 0.0000 | 0.0000 | 82 |
| pyramidal neuron | 0.6515 | 0.5947 | 0.6218 | 528 |

**Supplementary Table 10. Performance of native scGPT across cell types on the Zheng68k dataset**

| Cell type | Precision | Recall | F1-score | Support |
|---|---|---|---|---|
| CD4+ T Helper2 | 0.5000 | 0.0164 | 0.0317 | 61 |
| CD4+/CD25 T Reg | 0.4184 | 0.4044 | 0.4113 | 3672 |
| CD4+/CD45RA+/CD25- Naive T | 0.0809 | 0.0100 | 0.0179 | 1095 |
| CD4+/CD45RO+ Memory | 0.2305 | 0.1317 | 0.1676 | 1769 |
| CD8+ Cytotoxic T | 0.6454 | 0.5776 | 0.6096 | 12340 |
| CD8+/CD45RA+ Naive Cytotoxic | 0.5633 | 0.7787 | 0.6537 | 10161 |
| CD14+ Monocyte | 0.7516 | 0.8845 | 0.8127 | 1714 |
| CD19+ B | 0.9665 | 0.6626 | 0.7862 | 3524 |
| CD34+ | 1.0000 | 0.7972 | 0.8872 | 143 |
| CD56+ NK | 0.8332 | 0.9000 | 0.8653 | 5290 |
| Dendritic | 0.7544 | 0.5926 | 0.6638 | 1301 |

**Supplementary Table 11. Performance of native scGPT across cell types on the NSCLC dataset**

| Cell type | Precision | Recall | F1-score | Support |
|---|---|---|---|---|
| CD4_C1-Naive | 0.7135 | 0.7808 | 0.7456 | 3728 |
| CD4_C2-Tcm | 0.6230 | 0.6645 | 0.6431 | 1982 |
| CD4_C3-Tem | 0.6168 | 0.3376 | 0.4363 | 1205 |
| CD4_C4-CD69 | 0.6721 | 0.5114 | 0.5808 | 1539 |
| CD4_C5-ISG15 | 0.7656 | 0.8317 | 0.7973 | 1402 |
| CD4_C6-RPL | 0.7816 | 0.6228 | 0.6932 | 1034 |
| CD4_C7-Th1-like | 0.6071 | 0.2566 | 0.3607 | 1325 |
| CD4_C8-Treg | 0.7580 | 0.9214 | 0.8318 | 5714 |
| CD4_C9-Prolif. | 0.8684 | 0.8864 | 0.8773 | 484 |
| NA | 0.7262 | 0.5542 | 0.6286 | 4652 |
| Non-exhausted | 0.8417 | 0.9203 | 0.8792 | 11714 |
| Prolif. | 0.9238 | 0.8607 | 0.8911 | 732 |
| Tex | 0.7801 | 0.7976 | 0.7888 | 3132 |
| XCL1 | 0.0000 | 0.0000 | 0.0000 | 52 |

**Supplementary Table 12. Performance of native scGPT across cell types on the COVID dataset**

| Cell type | Precision | Recall | F1-score | Support |
|---|---|---|---|---|
| Activated CD4+ T | 0.8479 | 0.8668 | 0.8573 | 5075 |
| B | 0.9983 | 1.0000 | 0.9992 | 2955 |
| Effector Memory CD8+ T | 0.6263 | 0.7477 | 0.6817 | 547 |
| Effector T | 0.9159 | 0.8449 | 0.8790 | 13102 |
| Erythroid | 1.0000 | 0.8507 | 0.9194 | 67 |
| Innate T | 0.4103 | 0.7217 | 0.5232 | 1610 |
| Memory CD8+ T | 0.5535 | 0.3310 | 0.3982 | 849 |
| Mono DC | 0.9897 | 0.8075 | 0.8894 | 478 |
| Monocytes | 0.9866 | 0.9998 | 0.9932 | 5728 |
| NK | 0.8949 | 0.8948 | 0.8948 | 6214 |
| Naive CD4+ T | 0.8310 | 0.8067 | 0.8186 | 5705 |
| Naive CD8+ T | 0.6925 | 0.7656 | 0.7272 | 2338 |
| Plasma B | 0.9940 | 0.9940 | 0.9940 | 331 |
| Platelet | 1.0000 | 0.9978 | 0.9989 | 919 |
| Progenitor | 0.8910 | 0.8854 | 0.8882 | 157 |
| Proliferative T/NK | 0.9400 | 0.7091 | 0.8084 | 464 |

**Supplementary Table 13. Performance of full finetune scGPT across cell types on the MS dataset**

| Cell type | Precision | Recall | F1-score | Support |
|---|---|---|---|---|
| PVALB-expressing interneuron | 0.9485 | 0.9021 | 0.9247 | 878 |
| PVALB-expressing interneuron | 0.7796 | 0.9186 | 0.8434 | 258 |
| SV2C-expressing interneuron | 0.9477 | 0.8953 | 0.9208 | 344 |
| VIP-expressing interneuron | 0.9693 | 0.9810 | 0.9751 | 1158 |
| astrocyte | 0.9874 | 0.9406 | 0.9634 | 1330 |
| cortical layer 2-3 excitatory neuron A | 0.3101 | 0.5955 | 0.4079 | 314 |
| cortical layer 2-3 excitatory neuron B | 0.7672 | 0.8760 | 0.8180 | 2178 |
| cortical layer 4 excitatory neuron | 0.7998 | 0.8763 | 0.8363 | 1504 |
| cortical layer 5-6 excitatory neuron | 0.9130 | 0.6363 | 0.7499 | 1732 |
| endothelial cell | 0.9600 | 0.9882 | 0.9739 | 170 |
| microglial cell | 1.0000 | 0.3475 | 0.5157 | 118 |
| mixed excitatory neuron | 0.3153 | 0.1892 | 0.2365 | 185 |
| mixed glial cell | 0.7344 | 0.8394 | 0.7834 | 573 |
| oligodendrocyte A | 0.9474 | 0.9926 | 0.9695 | 1216 |
| oligodendrocyte C | 0.0000 | 0.0000 | 0.0000 | 34 |
| oligodendrocyte precursor cell | 0.9211 | 0.9711 | 0.9455 | 866 |
| phagocyte | 0.0000 | 0.0000 | 0.0000 | 82 |
| pyramidal neuron | 0.9335 | 0.7973 | 0.8601 | 528 |

**Supplementary Table 14. Performance of full finetune scGPT across cell types on the Zheng68k dataset**

| Cell type | Precision | Recall | F1-score | Support |
|---|---|---|---|---|
| CD4+ T Helper2 | 0.4167 | 0.0820 | 0.1370 | 61 |
| CD4+/CD25 T Reg | 0.6965 | 0.6236 | 0.6580 | 3672 |
| CD4+/CD45RA+/CD25- Naive T | 0.5096 | 0.2913 | 0.3707 | 1095 |
| CD4+/CD45RO+ Memory | 0.5285 | 0.4822 | 0.5043 | 1769 |
| CD8+ Cytotoxic T | 0.8457 | 0.8153 | 0.8303 | 12340 |
| CD8+/CD45RA+ Naive Cytotoxic | 0.7737 | 0.9188 | 0.8401 | 10161 |
| CD14+ Monocyte | 0.8342 | 0.8804 | 0.8567 | 1714 |
| CD19+ B | 0.8342 | 0.8804 | 0.8567 | 3524 |
| CD34+ | 0.9014 | 0.8951 | 0.8982 | 143 |
| CD56+ NK | 0.9258 | 0.9297 | 0.9277 | 5290 |
| Dendritic | 0.7843 | 0.7433 | 0.7632 | 1301 |

**Supplementary Table 15. Performance of full finetune scGPT across cell types on the NSCLC dataset**

| Cell type | Precision | Recall | F1-score | Support |
|---|---|---|---|---|
| CD4_C1-Naive | 0.8458 | 0.8594 | 0.8526 | 3728 |
| CD4_C2-Tcm | 0.8547 | 0.7240 | 0.7839 | 1982 |
| CD4_C3-Tem | 0.7718 | 0.6898 | 0.7285 | 1205 |
| CD4_C4-CD69 | 0.8416 | 0.8181 | 0.8297 | 1539 |
| CD4_C5-ISG15 | 0.9442 | 0.8452 | 0.8920 | 1402 |
| CD4_C6-RPL | 0.8518 | 0.7282 | 0.7852 | 1034 |
| CD4_C7-Th1-like | 0.9699 | 0.8506 | 0.9063 | 1325 |
| CD4_C8-Treg | 0.9009 | 0.9566 | 0.9279 | 5714 |
| CD4_C9-Prolif. | 0.9723 | 0.7975 | 0.8763 | 484 |
| NA | 0.7755 | 0.7999 | 0.7875 | 4652 |
| Non-exhausted | 0.9177 | 0.9318 | 0.9247 | 11714 |
| Prolif. | 0.9015 | 0.9003 | 0.9009 | 732 |
| Tex | 0.8283 | 0.9119 | 0.8681 | 3132 |
| XCL1 | 0.0000 | 0.0000 | 0.0000 | 52 |

**Supplementary Table 16. Performance of full finetune scGPT across cell types on the COVID dataset**

| Cell type | Precision | Recall | F1-score | Support |
|---|---|---|---|---|
| Activated CD4+ T | 0.9834 | 0.8390 | 0.9055 | 5075 |
| B | 0.9997 | 1.0000 | 0.9998 | 2955 |
| Effector Memory CD8+ T | 0.6502 | 0.9616 | 0.9616 | 547 |
| Effector T | 0.9451 | 0.9339 | 0.9395 | 13102 |
| Erythroid | 0.9828 | 0.8507 | 0.9120 | 67 |
| Innate T | 0.9212 | 0.8571 | 0.8880 | 1610 |
| Memory CD8+ T | 0.6282 | 0.7503 | 0.6838 | 849 |
| Mono DC | 0.9658 | 0.9456 | 0.9556 | 478 |
| Monocytes | 0.9956 | 0.9974 | 0.9965 | 5728 |
| NK | 0.9044 | 0.9121 | 0.9083 | 6214 |
| Naive CD4+ T | 0.9171 | 0.9770 | 0.9461 | 5705 |
| Naive CD8+ T | 0.9082 | 0.9855 | 0.9452 | 2338 |
| Plasma B | 0.9821 | 0.9970 | 0.9895 | 331 |
| Platelet | 1.0000 | 0.9967 | 0.9984 | 919 |
| Progenitor | 0.9671 | 0.9363 | 0.9515 | 157 |
| Proliferative T/NK | 0.8506 | 0.7974 | 0.8231 | 464 |

**Supplementary Table 17. Performance of finetune classifier scGPT across cell types on the MS dataset**

| Cell type | Precision | Recall | F1-score | Support |
|---|---|---|---|---|
| PVALB-expressing interneuron | 0.9323 | 0.7995 | 0.8608 | 878 |
| PVALB-expressing interneuron | 0.5898 | 0.8527 | 0.6973 | 258 |
| SV2C-expressing interneuron | 0.8775 | 0.8953 | 0.8863 | 344 |
| VIP-expressing interneuron | 0.9563 | 0.9456 | 0.9509 | 1158 |
| astrocyte | 0.9048 | 0.9714 | 0.9369 | 1330 |
| cortical layer 2-3 excitatory neuron A | 0.2332 | 0.5414 | 0.3260 | 314 |
| cortical layer 2-3 excitatory neuron B | 0.8740 | 0.6561 | 0.7495 | 2178 |
| cortical layer 4 excitatory neuron | 0.7317 | 0.8684 | 0.7942 | 1504 |
| cortical layer 5-6 excitatory neuron | 0.7207 | 0.7569 | 0.7384 | 1732 |
| endothelial cell | 0.8000 | 0.9882 | 0.8842 | 170 |
| microglial cell | 0.0000 | 0.0000 | 0.0000 | 118 |
| mixed excitatory neuron | 0.2632 | 0.0270 | 0.0490 | 185 |
| mixed glial cell | 0.7083 | 0.7627 | 0.7345 | 573 |
| oligodendrocyte A | 0.9446 | 0.9400 | 0.9423 | 1216 |
| oligodendrocyte C | 0.0000 | 0.0000 | 0.0000 | 34 |
| oligodendrocyte precursor cell | 0.9676 | 0.9319 | 0.9494 | 866 |
| phagocyte | 0.0000 | 0.0000 | 0.0000 | 82 |
| pyramidal neuron | 0.7196 | 0.7633 | 0.7408 | 528 |

**Supplementary Table 18. Performance of finetune classifier scGPT across cell types on the Zheng68k dataset**

| Cell type | Precision | Recall | F1-score | Support |
|---|---|---|---|---|
| CD4+ T Helper2 | 1.0000 | 0.0164 | 0.0323 | 61 |
| CD4+/CD25 T Reg | 0.4362 | 0.5664 | 0.4928 | 3672 |
| CD4+/CD45RA+/CD25- Naive T | 0.0000 | 0.0000 | 0.0000 | 1095 |
| CD4+/CD45RO+ Memory | 0.3741 | 0.0571 | 0.0991 | 1769 |
| CD8+ Cytotoxic T | 0.7626 | 0.5694 | 0.6520 | 12340 |
| CD8+/CD45RA+ Naive Cytotoxic | 0.5771 | 0.8722 | 0.6946 | 10161 |
| CD14+ Monocyte | 0.7401 | 0.9504 | 0.8322 | 1714 |
| CD19+ B | 0.9910 | 0.6564 | 0.7897 | 3524 |
| CD34+ | 1.0000 | 0.8392 | 0.9125 | 143 |
| CD56+ NK | 0.8360 | 0.9422 | 0.8859 | 5290 |
| Dendritic | 0.8304 | 0.5380 | 0.6530 | 1301 |

**Supplementary Table 19. Performance of finetune classifier scGPT across cell types on the NSCLC dataset**

| Cell type | Precision | Recall | F1-score | Support |
|---|---|---|---|---|
| CD4_C1-Naive | 0.7309 | 0.8415 | 0.7823 | 3728 |
| CD4_C2-Tcm | 0.7015 | 0.7043 | 0.7029 | 1982 |
| CD4_C3-Tem | 0.6584 | 0.4946 | 0.5649 | 1205 |
| CD4_C4-CD69 | 0.7315 | 0.6231 | 0.6730 | 1539 |
| CD4_C5-ISG15 | 0.7473 | 0.9051 | 0.8187 | 1402 |
| CD4_C6-RPL | 0.7572 | 0.7118 | 0.7338 | 1034 |
| CD4_C7-Th1-like | 0.6981 | 0.5638 | 0.6238 | 1325 |
| CD4_C8-Treg | 0.8260 | 0.9295 | 0.8747 | 5714 |
| CD4_C9-Prolif. | 0.9208 | 0.9132 | 0.9170 | 484 |
| NA | 0.7652 | 0.6369 | 0.6952 | 4652 |
| Non-exhausted | 0.9009 | 0.9165 | 0.9086 | 11714 |
| Prolif. | 0.9081 | 0.9180 | 0.9130 | 732 |
| Tex | 0.8644 | 0.8206 | 0.8419 | 3132 |
| XCL1 | 0.0000 | 0.0000 | 0.0000 | 52 |

**Supplementary Table 20. Performance of finetune classifier scGPT across cell types on the COVID dataset**

| Cell type | Precision | Recall | F1-score | Support |
|---|---|---|---|---|
| Activated CD4+ T | 0.9088 | 0.8703 | 0.8892 | 5075 |
| B | 1.0000 | 1.0000 | 1.0000 | 2955 |
| Effector Memory CD8+ T | 0.6702 | 0.9141 | 0.7734 | 547 |
| Effector T | 0.9184 | 0.9367 | 0.9275 | 13102 |
| Erythroid | 1.0000 | 0.8507 | 0.9194 | 67 |
| Innate T | 0.8134 | 0.7174 | 0.7624 | 1610 |
| Memory CD8+ T | 0.6639 | 0.4676 | 0.5487 | 849 |
| Mono DC | 0.9885 | 0.8954 | 0.9396 | 478 |
| Monocytes | 0.9924 | 0.9991 | 0.9957 | 5728 |
| NK | 0.9106 | 0.8964 | 0.9034 | 6214 |
| Naive CD4+ T | 0.8837 | 0.9229 | 0.9029 | 5705 |
| Naive CD8+ T | 0.8639 | 0.8961 | 0.8797 | 2338 |
| Plasma B | 0.9822 | 1.0000 | 0.9910 | 331 |
| Platelet | 1.0000 | 0.9989 | 0.9995 | 919 |
| Progenitor | 0.9236 | 0.9236 | 0.9236 | 157 |
| Proliferative T/NK | 0.9157 | 0.8427 | 0.8777 | 464 |

**Supplementary Table21. Performance of scGPT with Prefix prompt across cell types on the MS dataset**

| Cell type | Precision | Recall | F1-score | Support |
|---|---|---|---|---|
| PVALB-expressing interneuron | 0.9282 | 0.7654 | 0.8390 | 878 |
| PVALB-expressing interneuron | 0.5043 | 0.9109 | 0.6492 | 258 |
| SV2C-expressing interneuron | 0.8107 | 0.9215 | 0.8626 | 344 |
| VIP-expressing interneuron | 0.9681 | 0.8912 | 0.9281 | 1158 |
| astrocyte | 0.9653 | 0.9421 | 0.9536 | 1330 |
| cortical layer 2-3 excitatory neuron A | 0.2408 | 0.3535 | 0.2865 | 314 |
| cortical layer 2-3 excitatory neuron B | 0.8477 | 0.6850 | 0.7577 | 2178 |
| cortical layer 4 excitatory neuron | 0.7629 | 0.7467 | 0.7547 | 1504 |
| cortical layer 5-6 excitatory neuron | 0.6779 | 0.7182 | 0.6975 | 1732 |
| endothelial cell | 0.9441 | 0.9941 | 0.9685 | 170 |
| microglial cell | 0.9722 | 0.8898 | 0.9292 | 118 |
| mixed excitatory neuron | 0.1923 | 0.2162 | 0.2036 | 185 |
| mixed glial cell | 0.7274 | 0.8429 | 0.7809 | 573 |
| oligodendrocyte A | 0.9666 | 0.9507 | 0.9585 | 1216 |
| oligodendrocyte C | 0.2400 | 0.3529 | 0.2857 | 34 |
| oligodendrocyte precursor cell | 0.8992 | 0.9688 | 0.9327 | 866 |
| phagocyte | 0.6207 | 0.4390 | 0.5143 | 82 |
| pyramidal neuron | 0.6928 | 0.7860 | 0.7365 | 528 |

**Supplementary Table 22. Performance of scGPT with Prefix prompt across cell types on the Zheng68k dataset**

| Cell type | Precision | Recall | F1-score | Support |
|---|---|---|---|---|
| CD4+ T Helper2 | 0.0110 | 0.1148 | 0.0201 | 61 |
| CD4+/CD25 T Reg | 0.4493 | 0.3690 | 0.4052 | 3672 |
| CD4+/CD45RA+/CD25- Naive T | 0.0708 | 0.7616 | 0.1296 | 1095 |
| CD4+/CD45RO+ Memory | 0.1961 | 0.5777 | 0.2928 | 1769 |
| CD8+ Cytotoxic T | 0.9171 | 0.3622 | 0.5193 | 12340 |
| CD8+/CD45RA+ Naive Cytotoxic | 0.6768 | 0.2147 | 0.3260 | 10161 |
| CD14+ Monocyte | 0.7956 | 0.8629 | 0.8279 | 1714 |
| CD19+ B | 0.9532 | 0.6711 | 0.7877 | 3524 |
| CD34+ | 0.8742 | 0.9231 | 0.8980 | 143 |
| CD56+ NK | 0.7704 | 0.9773 | 0.8616 | 5290 |
| Dendritic | 0.7582 | 0.6580 | 0.7045 | 1301 |

**Supplementary Table 23. Performance of scGPT with Prefix prompt across cell types on the NSCLC dataset**

| Cell type | Precision | Recall | F1-score | Support |
|---|---|---|---|---|
| CD4_C1-Naive | 0.7815 | 0.7674 | 0.7744 | 3728 |
| CD4_C2-Tcm | 0.6090 | 0.7936 | 0.6892 | 1982 |
| CD4_C3-Tem | 0.4116 | 0.7356 | 0.5278 | 1205 |
| CD4_C4-CD69 | 0.6099 | 0.7264 | 0.6631 | 1539 |
| CD4_C5-ISG15 | 0.6740 | 0.9215 | 0.7785 | 1402 |
| CD4_C6-RPL | 0.6392 | 0.8172 | 0.7173 | 1034 |
| CD4_C7-Th1-like | 0.6070 | 0.7298 | 0.6628 | 1325 |
| CD4_C8-Treg | 0.9021 | 0.8276 | 0.8633 | 5714 |
| CD4_C9-Prolif. | 0.8366 | 0.9731 | 0.8997 | 484 |
| NA | 0.7338 | 0.5393 | 0.6217 | 4652 |
| Non-exhausted | 0.9611 | 0.7348 | 0.8329 | 11714 |
| Prolif. | 0.7505 | 0.9577 | 0.8415 | 732 |
| Tex | 0.7033 | 0.8902 | 0.7858 | 3132 |
| XCL1 | 0.0302 | 0.4038 | 0.0561 | 52 |

**Supplementary Table 24. Performance of scGPT with Prefix prompt across cell types on the COVID dataset**

| Cell type | Precision | Recall | F1-score | Support |
|---|---|---|---|---|
| Activated CD4+ T | 0.9237 | 0.7900 | 0.8516 | 5075 |
| B | 0.9993 | 0.9997 | 0.9995 | 2955 |
| Effector Memory CD8+ T | 0.6846 | 0.9324 | 0.7895 | 547 |
| Effector T | 0.9266 | 0.9318 | 0.9292 | 13102 |
| Erythroid | 0.8784 | 0.9701 | 0.9220 | 67 |
| Innate T | 0.7819 | 0.7839 | 0.7829 | 1610 |
| Memory CD8+ T | 0.4180 | 0.7350 | 0.5329 | 849 |
| Mono DC | 0.8528 | 0.9456 | 0.8968 | 478 |
| Monocytes | 0.9979 | 0.9860 | 0.9919 | 5728 |
| NK | 0.9304 | 0.8793 | 0.9041 | 6214 |
| Naive CD4+ T | 0.9173 | 0.8803 | 0.8984 | 5705 |
| Naive CD8+ T | 0.8356 | 0.9127 | 0.8724 | 2338 |
| Plasma B | 0.9763 | 0.9970 | 0.9865 | 331 |
| Platelet | 0.9989 | 0.9989 | 0.9989 | 919 |
| Progenitor | 0.8802 | 0.9363 | 0.9074 | 157 |
| Proliferative T/NK | 0.6976 | 0.9397 | 0.8007 | 464 |

**Supplementary Table 25. Performance of scGPT with LoRA prompt across cell types on the MS dataset**

| Cell type | Precision | Recall | F1-score | Support |
|---|---|---|---|---|
| PVALB-expressing interneuron | 0.9480 | 0.8519 | 0.8974 | 878 |
| PVALB-expressing interneuron | 0.6432 | 0.9225 | 0.7580 | 258 |
| SV2C-expressing interneuron | 0.7901 | 0.9738 | 0.8724 | 344 |
| VIP-expressing interneuron | 0.9771 | 0.9231 | 0.9494 | 1158 |
| astrocyte | 0.9792 | 0.9534 | 0.9661 | 1330 |
| cortical layer 2-3 excitatory neuron A | 0.3500 | 0.4459 | 0.3922 | 314 |
| cortical layer 2-3 excitatory neuron B | 0.8578 | 0.8058 | 0.8310 | 2178 |
| cortical layer 4 excitatory neuron | 0.7866 | 0.8604 | 0.8218 | 1504 |
| cortical layer 5-6 excitatory neuron | 0.8815 | 0.6917 | 0.7752 | 1732 |
| endothelial cell | 0.9767 | 0.9882 | 0.9825 | 170 |
| microglial cell | 0.9626 | 0.8729 | 0.9156 | 118 |
| mixed excitatory neuron | 0.1503 | 0.2649 | 0.1918 | 185 |
| mixed glial cell | 0.7652 | 0.8586 | 0.8092 | 573 |
| oligodendrocyte A | 0.9677 | 0.9613 | 0.9645 | 1216 |
| oligodendrocyte C | 0.1918 | 0.4118 | 0.2617 | 34 |
| oligodendrocyte precursor cell | 0.9225 | 0.9758 | 0.9484 | 866 |
| phagocyte | 0.6452 | 0.4878 | 0.5556 | 82 |
| pyramidal neuron | 0.7106 | 0.7254 | 0.7179 | 528 |

**Supplementary Table 26. Performance of scGPT with LoRA prompt across cell types on the Zheng68k dataset**

| Cell type | Precision | Recall | F1-score | Support |
|---|---|---|---|---|
| CD4+ T Helper2 | 0.0166 | 0.1148 | 0.0290 | 61 |
| CD4+/CD25 T Reg | 0.5123 | 0.4153 | 0.4587 | 3672 |
| CD4+/CD45RA+/CD25- Naive T | 0.0889 | 0.6374 | 0.1561 | 1095 |
| CD4+/CD45RO+ Memory | 0.2596 | 0.6009 | 0.3626 | 1769 |
| CD8+ Cytotoxic T | 0.8704 | 0.5370 | 0.6642 | 12340 |
| CD8+/CD45RA+ Naive Cytotoxic | 0.6777 | 0.4365 | 0.5310 | 10161 |
| CD14+ Monocyte | 0.8623 | 0.6908 | 0.7671 | 1714 |
| CD19+ B | 0.9220 | 0.7046 | 0.7988 | 3524 |
| CD34+ | 0.7778 | 0.9301 | 0.8471 | 143 |
| CD56+ NK | 0.8599 | 0.9302 | 0.8937 | 5290 |
| Dendritic | 0.6297 | 0.7802 | 0.6969 | 1301 |

**Supplementary Table 27. Performance of scGPT with LoRA prompt across cell types on the NSCLC dataset**

| Cell type | Precision | Recall | F1-score | Support |
|---|---|---|---|---|
| CD4_C1-Naive | 0.7050 | 0.8391 | 0.7662 | 3728 |
| CD4_C2-Tcm | 0.6140 | 0.7992 | 0.6944 | 1982 |
| CD4_C3-Tem | 0.5166 | 0.6234 | 0.5650 | 1205 |
| CD4_C4-CD69 | 0.6781 | 0.7089 | 0.6931 | 1539 |
| CD4_C5-ISG15 | 0.8390 | 0.8509 | 0.8449 | 1402 |
| CD4_C6-RPL | 0.5427 | 0.8859 | 0.6730 | 1034 |
| CD4_C7-Th1-like | 0.6295 | 0.8936 | 0.7386 | 1325 |
| CD4_C8-Treg | 0.9365 | 0.7998 | 0.8628 | 5714 |
| CD4_C9-Prolif. | 0.8981 | 0.9649 | 0.9303 | 484 |
| NA | 0.6705 | 0.5464 | 0.6022 | 4652 |
| Non-exhausted | 0.9589 | 0.6137 | 0.7484 | 11714 |
| Prolif. | 0.9181 | 0.8880 | 0.9028 | 732 |
| Tex | 0.4935 | 0.9460 | 0.6486 | 3132 |
| XCL1 | 0.1341 | 0.6731 | 0.2236 | 52 |

**Supplementary Table 28. Performance of scGPT with LoRA prompt across cell types on the COVID dataset**

| Cell type | Precision | Recall | F1-score | Support |
|---|---|---|---|---|
| Activated CD4+ T | 0.9813 | 0.7742 | 0.8655 | 5075 |
| B | 0.9986 | 1.0000 | 0.9993 | 2955 |
| Effector Memory CD8+ T | 0.8913 | 0.8245 | 0.8566 | 547 |
| Effector T | 0.9707 | 0.8627 | 0.9135 | 13102 |
| Erythroid | 0.9041 | 0.9851 | 0.9429 | 67 |
| Innate T | 0.6537 | 0.9602 | 0.7779 | 1610 |
| Memory CD8+ T | 0.4611 | 0.8445 | 0.5965 | 849 |
| Mono DC | 0.7308 | 0.9770 | 0.8362 | 478 |
| Monocytes | 0.9996 | 0.9689 | 0.9840 | 5728 |
| NK | 0.8838 | 0.9207 | 0.9019 | 6214 |
| Naive CD4+ T | 0.8793 | 0.9706 | 0.9227 | 5705 |
| Naive CD8+ T | 0.9149 | 0.9748 | 0.9439 | 2338 |
| Plasma B | 0.9851 | 0.9970 | 0.9910 | 331 |
| Platelet | 0.9989 | 1.0000 | 0.9995 | 919 |
| Progenitor | 0.9304 | 0.9363 | 0.9333 | 157 |
| Proliferative T/NK | 0.7729 | 0.9461 | 0.8508 | 464 |

**Supplementary Table 29. Performance of scGPT with Gene token prompt across cell types on the MS dataset**

| Cell type | Precision | Recall | F1-score | Support |
|---|---|---|---|---|
| PVALB-expressing interneuron | 0.9538 | 0.8702 | 0.9101 | 878 |
| PVALB-expressing interneuron | 0.6997 | 0.9031 | 0.7885 | 258 |
| SV2C-expressing interneuron | 0.8247 | 0.9709 | 0.8919 | 344 |
| VIP-expressing interneuron | 0.9792 | 0.9370 | 0.9576 | 1158 |
| astrocyte | 0.9798 | 0.9496 | 0.9645 | 1330 |
| cortical layer 2-3 excitatory neuron A | 0.3052 | 0.7484 | 0.4336 | 314 |
| cortical layer 2-3 excitatory neuron B | 0.9118 | 0.7075 | 0.7968 | 2178 |
| cortical layer 4 excitatory neuron | 0.8342 | 0.8231 | 0.8286 | 1504 |
| cortical layer 5-6 excitatory neuron | 0.8727 | 0.7523 | 0.8081 | 1732 |
| endothelial cell | 0.9337 | 0.9941 | 0.9630 | 170 |
| microglial cell | 0.9717 | 0.8729 | 0.9196 | 118 |
| mixed excitatory neuron | 0.1561 | 0.2919 | 0.2034 | 185 |
| mixed glial cell | 0.7489 | 0.8901 | 0.8134 | 573 |
| oligodendrocyte A | 0.9742 | 0.9622 | 0.9681 | 1216 |
| oligodendrocyte C | 0.2812 | 0.2647 | 0.2727 | 34 |
| oligodendrocyte precursor cell | 0.9422 | 0.9792 | 0.9604 | 866 |
| phagocyte | 0.7358 | 0.4756 | 0.5778 | 82 |
| pyramidal neuron | 0.7378 | 0.8314 | 0.7818 | 528 |

**Supplementary Table 30. Performance of scGPT with Gene token prompt across cell types on the Zheng68k dataset**

| Cell type | Precision | Recall | F1-score | Support |
|---|---|---|---|---|
| CD4+ T Helper2 | 0.0690 | 0.1311 | 0.0904 | 61 |
| CD4+/CD25 T Reg | 0.5311 | 0.6155 | 0.5702 | 3672 |
| CD4+/CD45RA+/CD25- Naive T | 0.1390 | 0.5963 | 0.2255 | 1095 |
| CD4+/CD45RO+ Memory | 0.2647 | 0.7168 | 0.3866 | 1769 |
| CD8+ Cytotoxic T | 0.9264 | 0.4981 | 0.6478 | 12340 |
| CD8+/CD45RA+ Naive Cytotoxic | 0.7733 | 0.5573 | 0.6478 | 10161 |
| CD14+ Monocyte | 0.8045 | 0.8909 | 0.8455 | 1714 |
| CD19+ B | 0.8376 | 0.8930 | 0.8644 | 3524 |
| CD34+ | 0.8487 | 0.9021 | 0.8746 | 143 |
| CD56+ NK | 0.8193 | 0.9745 | 0.8902 | 5290 |
| Dendritic | 0.7706 | 0.6841 | 0.7248 | 1301 |

**Supplementary Table 31. Performance of scGPT with Gene token prompt across cell types on the NSCLC dataset**

| Cell type | Precision | Recall | F1-score | Support |
|---|---|---|---|---|
| CD4_C1-Naive | 0.8842 | 0.8294 | 0.8559 | 3728 |
| CD4_C2-Tcm | 0.7470 | 0.8910 | 0.8127 | 1982 |
| CD4_C3-Tem | 0.6570 | 0.8185 | 0.7289 | 1205 |
| CD4_C4-CD69 | 0.7898 | 0.8934 | 0.8384 | 1539 |
| CD4_C5-ISG15 | 0.8351 | 0.9608 | 0.8935 | 1402 |
| CD4_C6-RPL | 0.7605 | 0.8443 | 0.8002 | 1034 |
| CD4_C7-Th1-like | 0.9046 | 0.9449 | 0.9243 | 1325 |
| CD4_C8-Treg | 0.9487 | 0.9067 | 0.9272 | 5714 |
| CD4_C9-Prolif. | 0.9055 | 0.9504 | 0.9274 | 484 |
| NA | 0.8078 | 0.7794 | 0.7933 | 4652 |
| Non-exhausted | 0.9626 | 0.855 | 0.9059 | 11714 |
| Prolif. | 0.8422 | 0.9549 | 0.8950 | 732 |
| Tex | 0.7930 | 0.9246 | 0.8538 | 3132 |
| XCL1 | 0.3121 | 0.8462 | 0.4560 | 52 |

**Supplementary Table 32. Performance of scGPT with Gene token prompt across cell types on the COVID dataset**

| Cell type | Precision | Recall | F1-score | Support |
|---|---|---|---|---|
| Activated CD4+ T | 0.9253 | 0.9302 | 0.9278 | 5075 |
| B | 0.9997 | 1.0000 | 0.9998 | 2955 |
| Effector Memory CD8+ T | 0.8204 | 0.9104 | 0.8631 | 547 |
| Effector T | 0.9691 | 0.9257 | 0.9469 | 13102 |
| Erythroid | 0.8049 | 0.9851 | 0.8859 | 67 |
| Innate T | 0.8535 | 0.9590 | 0.9032 | 1610 |
| Memory CD8+ T | 0.7255 | 0.8622 | 0.7879 | 849 |
| Mono DC | 0.9201 | 0.9393 | 0.9296 | 478 |
| Monocytes | 0.9981 | 0.9930 | 0.9955 | 5728 |
| NK | 0.8897 | 0.9329 | 0.9108 | 6214 |
| Naive CD4+ T | 0.9695 | 0.9246 | 0.9465 | 5705 |
| Naive CD8+ T | 0.9534 | 0.9705 | 0.9618 | 2338 |
| Plasma B | 0.9821 | 0.9970 | 0.9895 | 331 |
| Platelet | 1.0000 | 1.0000 | 1.0000 | 919 |
| Progenitor | 0.8802 | 0.9363 | 0.9074 | 157 |
| Proliferative T/NK | 0.8382 | 0.9267 | 0.8802 | 464 |

**Supplementary Table 33. Performance of scGPT with Gene encoder prompt across cell types on the MS dataset**

| Cell type | Precision | Recall | F1-score | Support |
|---|---|---|---|---|
| PVALB-expressing interneuron | 0.9396 | 0.9214 | 0.9304 | 878 |
| PVALB-expressing interneuron | 0.7857 | 0.8527 | 0.8178 | 258 |
| SV2C-expressing interneuron | 0.8616 | 0.9593 | 0.9078 | 344 |
| VIP-expressing interneuron | 0.9796 | 0.9525 | 0.9658 | 1158 |
| astrocyte | 0.9841 | 0.9774 | 0.9808 | 1330 |
| cortical layer 2-3 excitatory neuron A | 0.3739 | 0.6752 | 0.4813 | 314 |
| cortical layer 2-3 excitatory neuron B | 0.8958 | 0.7975 | 0.8438 | 2178 |
| cortical layer 4 excitatory neuron | 0.8220 | 0.8564 | 0.8388 | 1504 |
| cortical layer 5-6 excitatory neuron | 0.8771 | 0.7748 | 0.8228 | 1732 |
| endothelial cell | 0.9766 | 0.9824 | 0.9795 | 170 |
| microglial cell | 0.9633 | 0.8898 | 0.9251 | 118 |
| mixed excitatory neuron | 0.2266 | 0.3405 | 0.2721 | 185 |
| mixed glial cell | 0.8142 | 0.9023 | 0.8560 | 573 |
| oligodendrocyte A | 0.9666 | 0.9762 | 0.9714 | 1216 |
| oligodendrocyte C | 0.5000 | 0.2941 | 0.3704 | 34 |
| oligodendrocyte precursor cell | 0.9581 | 0.9769 | 0.9674 | 866 |
| phagocyte | 0.8000 | 0.3415 | 0.4786 | 82 |
| pyramidal neuron | 0.8355 | 0.8466 | 0.8410 | 528 |

**Supplementary Table 34. Performance of scGPT with Gene encoder prompt across cell types on the Zheng68k dataset**

| Cell type | Precision | Recall | F1-score | Support |
|---|---|---|---|---|
| CD4+ T Helper2 | 0.0714 | 0.1311 | 0.0925 | 61 |
| CD4+/CD25 T Reg | 0.5501 | 0.6887 | 0.6117 | 3672 |
| CD4+/CD45RA+/CD25- Naive T | 0.1444 | 0.6530 | 0.2365 | 1095 |
| CD4+/CD45RO+ Memory | 0.3164 | 0.6789 | 0.4316 | 1769 |
| CD8+ Cytotoxic T | 0.9361 | 0.5593 | 0.7002 | 12340 |
| CD8+/CD45RA+ Naive Cytotoxic | 0.8084 | 0.5653 | 0.6654 | 10161 |
| CD14+ Monocyte | 0.8076 | 0.9160 | 0.8584 | 1714 |
| CD19+ B | 0.9011 | 0.8788 | 0.8898 | 3524 |
| CD34+ | 0.9085 | 0.9021 | 0.9053 | 143 |
| CD56+ NK | 0.8080 | 0.9875 | 0.8888 | 5290 |
| Dendritic | 0.7831 | 0.6910 | 0.7342 | 1301 |

**Supplementary Table 35. Performance of scGPT with Gene encoder prompt across cell types on the NSCLC dataset**

| Cell type | Precision | Recall | F1-score | Support |
|---|---|---|---|---|
| CD4_C1-Naive | 0.9086 | 0.8028 | 0.8525 | 3728 |
| CD4_C2-Tcm | 0.7654 | 0.8789 | 0.8182 | 1982 |
| CD4_C3-Tem | 0.5781 | 0.8556 | 0.6900 | 1205 |
| CD4_C4-CD69 | 0.7720 | 0.8824 | 0.8235 | 1539 |
| CD4_C5-ISG15 | 0.8233 | 0.9672 | 0.8895 | 1402 |
| CD4_C6-RPL | 0.7134 | 0.8762 | 0.7865 | 1034 |
| CD4_C7-Th1-like | 0.8921 | 0.9608 | 0.9251 | 1325 |
| CD4_C8-Treg | 0.9449 | 0.9121 | 0.9282 | 5714 |
| CD4_C9-Prolif. | 0.9111 | 0.9525 | 0.9313 | 484 |
| NA | 0.8523 | 0.7369 | 0.7904 | 4652 |
| Non-exhausted | 0.9539 | 0.8943 | 0.9232 | 11714 |
| Prolif. | 0.8423 | 0.9631 | 0.8987 | 732 |
| Tex | 0.8535 | 0.9112 | 0.8814 | 3132 |
| XCL1 | 0.3390 | 0.7692 | 0.4706 | 52 |

**Supplementary Table 36. Performance of scGPT with Gene encoder prompt across cell types on the COVID dataset**

| Cell type | Precision | Recall | F1-score | Support |
|---|---|---|---|---|
| Activated CD4+ T | 0.9296 | 0.9310 | 0.9303 | 5075 |
| B | 0.9997 | 1.0000 | 0.9998 | 2955 |
| Effector Memory CD8+ T | 0.8426 | 0.9397 | 0.8885 | 547 |
| Effector T | 0.9720 | 0.9285 | 0.9497 | 13102 |
| Erythroid | 0.7976 | 1.0000 | 0.8874 | 67 |
| Innate T | 0.8608 | 0.9602 | 0.9078 | 1610 |
| Memory CD8+ T | 0.6831 | 0.9117 | 0.7810 | 849 |
| Mono DC | 0.8844 | 0.9603 | 0.9208 | 478 |
| Monocytes | 0.9977 | 0.9894 | 0.9935 | 5728 |
| NK | 0.8968 | 0.9316 | 0.9139 | 6214 |
| Naive CD4+ T | 0.9759 | 0.9280 | 0.9513 | 5705 |
| Naive CD8+ T | 0.9621 | 0.9675 | 0.9648 | 2338 |
| Plasma B | 0.9881 | 1.0000 | 0.9940 | 331 |
| Platelet | 0.9989 | 1.0000 | 0.9995 | 919 |
| Progenitor | 0.9363 | 0.9363 | 0.9363 | 157 |
| Proliferative T/NK | 0.8227 | 0.9203 | 0.8688 | 464 |

**Supplementary Table 37. Performance of different hyper parameters in Prefix prompt**

| Dataset | Number of tokens | Accuracy | Precision | Recall | F1-score |
|---|---|---|---|---|---|
| M.S. | 16 | 0.721 | 0.715 | 0.710 | 0.732 |
| | 32 | 0.742 | 0.781 | 0.704 | 0.761 |
| | 64 | **0.743** | **0.815** | **0.797** | **0.802** |
| | 128 | 0.724 | 0.820 | 0.764 | 0.780 |
| | 256 | 0.704 | 0.733 | 0.685 | 0.707 |
| Zheng68k | 16 | 0.506 | 0.670 | 0.485 | 0.504 |
| | 32 | 0.571 | 0.722 | 0.463 | 0.521 |
| | 64 | **0.590** | **0.735** | 0.484 | **0.528** |
| | 128 | 0.580 | 0.733 | 0.483 | 0.522 |
| | 256 | 0.565 | 0.618 | **0.540** | 0.509 |
| NSCLC | 16 | 0.471 | 0.480 | 0.445 | 0.460 |
| | 32 | 0.588 | 0.580 | 0.474 | 0.482 |
| | 64 | **0.590** | **0.735** | 0.484 | **0.528** |
| | 128 | **0.590** | 0.521 | 0.460 | 0.515 |
| | 256 | 0.567 | 0.573 | **0.542** | 0.458 |
| COVID | 16 | 0.891 | 0.903 | 0.885 | 0.894 |
| | 32 | 0.906 | 0.912 | 0.894 | 0.906 |
| | 64 | **0.914** | **0.916** | **0.906** | **0.909** |
| | 128 | 0.898 | 0.902 | 0.890 | 0.901 |
| | 256 | 0.886 | 0.889 | 0.867 | 0.880 |

**Supplementary Table 38. Performance of different hyper parameters in LoRA prompt**

| Dataset | Method | Accuracy | Precision | Recall | F1-score |
|---|---|---|---|---|---|
| ms | Rank-8 Alpha-1 | **0.787** | **0.873** | **0.875** | **0.847** |
| | Rank-8 Alpha-8 | 0.774 | 0.780 | 0.779 | 0.832 |
| | Rank-64 Alpha-8 | 0.775 | 0.812 | 0.811 | 0.846 |
| | Rank-64 Alpha-128 | 0.739 | 0.808 | 0.833 | 0.765 |
| COVID | Rank-8 Alpha-1 | **0.939** | **0.941** | **0.944** | **0.925** |
| | Rank-8 Alpha-8 | 0.904 | 0.905 | 0.922 | 0.910 |
| | Rank-64 Alpha-8 | 0.911 | 0.897 | 0.891 | 0.899 |
| | Rank-64 Alpha-128 | 0.865 | 0.899 | 0.830 | 0.870 |
| Zheng68k | Rank-8 Alpha-1 | 0.599 | **0.771** | **0.673** | 0.630 |
| | Rank-8 Alpha-8 | 0.568 | 0.612 | 0.522 | 0.661 |
| | Rank-64 Alpha-8 | **0.600** | 0.698 | 0.656 | 0.655 |
| | Rank-64 Alpha-128 | 0.552 | 0.605 | 0.518 | **0.672** |
| NSCLC | Rank-8 Alpha-1 | 0.715 | 0.870 | **0.824** | 0.737 |
| | Rank-8 Alpha-8 | **0.772** | 0.753 | 0.729 | 0.653 |
| | Rank-64 Alpha-8 | 0.716 | 0.804 | 0.801 | 0.694 |
| | Rank-64 Alpha-128 | 0.726 | **0.894** | 0.791 | **0.757** |

**Supplementary Table 39. Performance of different hyper parameters in Gene encoder prompt**

| Dataset | Method | Accuracy | Precision | Recall | F1-score |
|---|---|---|---|---|---|
| ms | Ahead-6 | **0.795** | **0.884** | **0.870** | **0.874** |
| | Behind-6 | 0.701 | 0.763 | 0.770 | 0.823 |
| | Ahead-3 | 0.745 | 0.823 | 0.821 | 0.848 |
| | Behind-3 | 0.757 | 0.835 | 0.822 | 0.847 |
| | Ahead-1 | 0.754 | 0.834 | 0.812 | 0.846 |
| | Ahead-2 | 0.777 | 0.846 | 0.825 | 0.844 |
| | Ahead-3 | 0.759 | 0.838 | 0.829 | 0.855 |
| COVID | Ahead-6 | **0.957** | 0.949 | **0.946** | **0.947** |
| | Behind-6 | 0.902 | 0.921 | 0.930 | 0.920 |
| | Ahead-3 | 0.931 | 0.947 | 0.940 | 0.941 |
| | Behind-3 | 0.944 | 0.932 | 0.934 | 0.944 |
| | Ahead-1 | 0.951 | 0.948 | 0.945 | 0.934 |
| | Ahead-2 | 0.945 | 0.954 | 0.923 | 0.921 |
| | Ahead-3 | 0.949 | **0.955** | 0.943 | 0.934 |
| Zheng68k | Ahead-6 | **0.696** | **0.791** | **0.682** | **0.708** |
| | Behind-6 | 0.568 | 0.605 | 0.547 | 0.614 |
| | Ahead-3 | 0.623 | 0.678 | 0.615 | 0.654 |
| | Behind-3 | 0.631 | 0.702 | 0.613 | 0.660 |
| | Ahead-1 | 0.633 | 0.693 | 0.620 | 0.649 |
| | Ahead-2 | 0.637 | 0.706 | 0.602 | 0.653 |
| | Ahead-3 | 0.616 | 0.705 | 0.598 | 0.662 |
| NSCLC | Ahead-6 | **0.883** | **0.884** | **0.874** | **0.876** |
| | Behind-6 | 0.772 | 0.762 | 0.738 | 0.743 |
| | Ahead-3 | 0.833 | 0.834 | 0.811 | 0.823 |
| | Behind-3 | 0.821 | 0.827 | 0.808 | 0.809 |
| | Ahead-1 | 0.835 | 0.816 | 0.810 | 0.814 |
| | Ahead-2 | 0.828 | 0.814 | 0.817 | 0.802 |
| | Ahead-3 | 0.838 | 0.827 | 0.799 | 0.809 |
